# Stingless bees, turtle ants and tea plants are rich sources of undescribed *Lactobacillaceae*

**DOI:** 10.64898/2026.08.12.744358

**Authors:** Scott A. Oliphant, Jennifer M. Gardner, Vladimir Jiranek, Krista M. Sumby

## Abstract

Knowledge of the family *Lactobacillaceae* rests largely on isolates from foods and a few repeatedly sampled hosts. Reference databases give an unrecognised sequence the name of its nearest relative, so a lineage with no entry of its own is renamed rather than flagged. Here we classify the family across the public plant and invertebrate amplicon record to locate the hosts carrying undescribed lineages. Across 3,344 independent 16S rRNA gene amplicon studies, every sequence cluster was tested against a type-strain reference and placed at the deepest rank it supports. Of 204,813 classified clusters, 20,516 were named to species and 127,906 to genus, while 4,741 matched no described species. Six published datasets whose authors could name their *Lactobacillaceae* only as “*Lactobacillus*”, or not at all, are reclassified here. Genera described from one habitat occur far beyond it, three bee-associated genera occurring on Rosaceae and Brassicaceae at several times their rate on wind-pollinated grasses, and we found no published *Bombilactobacillus* record from a plant. Undescribed lineages concentrate in the least-cultured hosts, reaching 86.1% and 77.4% of studies in the stingless bees *Melipona* and *Tetragonula*, and are most divergent in the turtle ant *Cephalotes* and, among plants, in tea, *Camellia*. An independent genome-resolved survey of pot honey converges on the same two genera. The primary descriptions of forty-eight species from these hosts specify a supplemented medium, so the hosts carrying undescribed lineages also indicate how to culture them.

## INTRODUCTION

Lactic acid bacteria are Gram-positive, fermentative microorganisms recovered from food, plant, vertebrate-gut and invertebrate niches. Their evolutionary history has been interpreted as involving transitions among free-living, nomadic and host-adapted lifestyles [1]. The family *Lactobacillaceae* is the taxonomic core of the group and, since the 2020 union with *Leuconostocaceae*, sits in one nomenclatural framework holding 467 validly published type-strain entries, 441 species and 26 subspecies across 37 genera [2–4].

Within insects, host adaptation in this family is common. Honey bees carry *Apilactobacillus*, *Bombilactobacillus* and *Lactobacillus* consistently wherever they are sampled, and flowers, fruits and fermented plant material supply the fructophilic species found alongside them [5–8]. That specificity extends to few host lineages. Only honey bees and bumble bees harbour lactobacilli of their own, while sweat bees carry relatives of flower-inhabiting fructophiles and ants carry relatives of vertebrate and environmental strains [9]. However, an isolation record does not necessarily identify a microorganism’s natural habitat. Lactobacilli may be allochthonous, with no ecological or evolutionary connection to the habitat they are recovered from [1]. Most knowledge of the family still derives from human-made systems, so its ecological context in nature remains poorly understood [10]. Recent taxonomic work has likewise focused on a narrow range of hosts, with five *Apilactobacillus* and *Bombilactobacillus* species described from *Apis* guts and honey between 2021 and 2026 alone [11–14].

Reference databases compound this problem by assigning queries to the closest labelled entry available. A lineage with no entry of its own is therefore given the name of a relative, usually without warning, and seven genera of this family carry no label at all in SILVA 138.2. Published work shows the consequence. Ramalho and Moreau [15] could assign the *Lactobacillaceae* in the core gut community of more than 75 *Cephalotes* species only to “*Lactobacillus*”, while Mills *et al*. [16] identified *Bombilactobacillus thymidiniphilus* in *Tetragonula carbonaria* only by searching outside the database with BLAST. In both cases, a described member of the family was present, but the database could not resolve the sequence to it.

However, a full-length sequence exists for almost every type strain of this family [4]. Against this reference, Unassigner estimates the probability that a query’s full-length identity to a described species falls below the species boundary, using only the query’s own amplicon window [17]. Every described species can therefore be tested in turn: a query surviving against several species of one genus is placed at genus, and one ruled out of everything is treated as undescribed rather than renamed.

Here we apply that rule across the public plant and invertebrate 16S rRNA gene amplicon record, comprising 3,344 independent BioProjects. We recover named genera and species in datasets whose authors could not name them, identify the hosts carrying the deepest undescribed lineages, and show from the type-strain cultivation record that many of those lineages will require supplementation to grow. Parente *et al*. [18] surveyed this family across foods and food environments. This study surveys it across the plant and invertebrate hosts.

## MATERIALS AND METHODS

### Cultivation and media record

Cultivation and media records were compiled from the original description of each species’ nomenclatural type strain. Nine genera in which at least 60% of described species have an insect or plant isolation source were surveyed, along with exemplar species from larger mixed genera. The type strain’s host, the medium recipe as stated, the fructophilic type, and the strain’s BacDive and MediaDive record were recorded for each species whose description reports a medium supplement (Table S1). Further details, including the reference phylogeny shown in Figure S1, are provided in the Supplementary Methods.

### Amplicon dataset

Public 16S rRNA gene amplicon runs were retrieved from the NCBI Sequence Read Archive in acquisition waves scoped by NCBI taxonomy and restricted at run level to plant and invertebrate hosts, the analysed scope being the first four waves (Supplementary Methods). 948,386 runs were processed across the sweep, and the 3,284 BioProjects in which the family was resolved to genus or species rank are the denominator of every fitted quantity (Table S2). Host was resolved with TaxonKit [19] against a pinned NCBI taxonomy dump from four metadata fields, in order: the BioSample host scientific name, the BioSample `host` attribute, the run title and the isolation source. The rules for an unresolved or a contradicted host are in Supplementary Methods. Control, mock and blank runs were identified from sample metadata, excluded from every aggregation, and used as the independent panel for the contamination screen below (Table S3a).

Runs were downloaded with the SRA Toolkit [20, 21] and the generic Illumina 3’ adapter removed with Cutadapt [22]. The remaining steps used VSEARCH [23]. Paired reads were merged where they overlapped, quality-filtered [24] and dereplicated strand-aware with within-sample singletons removed [25]. Where fewer than one read in five merged, the forward read was carried alone. Primers were not trimmed. Reads were clustered into fine-resolution OTUs with Swarm at *d* = 1 in fastidious mode [26–28]. The OTU is the unit reported throughout and its seed sequence was the query classified. Seeds were screened for chimeras [29], and chimera-free seeds were recruited against a curated *Lactobacillaceae* 16S rRNA gene reference at 80% identity [30], below the 86.5% family threshold of Yarza *et al*. [31]. Recruitment decides only whether a read is plausibly in the family and uses a different reference from the rank calls (Supplementary Methods). Recruited reads were pooled, dereplicated across runs, and reduced to those OTUs recurring in at least two independent BioProjects. The amplicon region of each surviving OTU was read from its own sequence against the *Escherichia coli* coordinate frame [32]. Every tool, version and non-default parameter is in Table S4, and the effect of each choice, including the benchmark preferring Swarm to denoising [33], is in Supplementary Methods.

### Classification

OTUs were classified in three stages against LactoTypeDB v1.0.0, our type-anchored 16S rRNA gene reference for the family, in the order that reference recommends (Supplementary Methods) [4]. Family membership was assigned with DADA2 `assignTaxonomy` [34], which implements the naive Bayesian classifier of Wang *et al*. [35], against SILVA 138.2 SSU Ref NR99 unmodified. Each in-family OTU was then tested against individual species with Unassigner [17] against LactoTypeDB at its hard default. An OTU is ruled out of a type strain where the probability that full-length identity falls below 97.5% reaches 0.5, and the species that survive this test form the compatible set. The third stage applies artefact screens to OTUs returning an empty set, removing chimeric, borderline-chimeric, multi-copy *rrn* and non-type conspecific ones (Supplementary Methods).

The rank assignment is specific to this study. Each in-family OTU was placed at the deepest rank its own evidence supports, its compatible set first resolved to canonical species through the reference’s accession-to-species table. A set holding one species was named at species, a set spanning several species of one genus at genus, and a non-empty set spanning more than one genus at family rank, termed ambiguous. An OTU with an empty set was assigned against three identity thresholds. At or above 98.65% [36] an empty set contradicts the OTU’s own identity to a described species, and the OTU is dropped from the reported detections. Between 98.65 and 94.5% [31] it is treated as an undescribed species, and between 94.5% and the 86.5% family threshold as an undescribed genus. All three are used as reference points rather than as decision rules (Supplementary Methods). The two undescribed classes are reported as one category, with identity to the nearest type strain reported as a continuous quantity.

An undescribed OTU has no genus assignment by definition, so it was assigned to the genus of its nearest described type strain, but only where that type strain is the sole match at the OTU’s own V region or falls in a collapse group containing a single genus. In every other case the OTU is left unassigned, and the unassigned set is reported as its own quantity and never distributed across genera.

### Controls

For each genus, the OTU seed sequences carrying its analysed detections were compared with those in the negative control runs, which were excluded from the analysed dataset. The fraction of the analysed set also occurring in a control is reported (Table S3b). Control panel definitions and the separation of reagent background from plate carryover are in Supplementary Methods.

Four further controls are reported in Supplementary Methods: recovery of a published lineage from an independent BioProject, rejection of out-of-family sequences at the family-membership test, completeness of the reference, and the behaviour of that test with parts of the reference withheld.

### Statistical analysis

The unit of analysis is one BioProject at one host node, a host node being a host taxon at order, family or genus rank or the host class itself, with the rank reported on every estimate. Host class takes two values, plant and invertebrate. Only units in which the family was detected were analysed. Runs deposited more than once, and BioProject accessions identified as mirrors of one dataset, were removed before any aggregation (Tables S5a, S5b).

Two quantities were fitted the same way: the probability that a genus is detected in a BioProject of a particular host node, conditional on the family being detected there, and the probability that a BioProject yields an undescribed OTU. Both were fitted as `outcome` `∼` `node` + `z(log10 depth) + z(log10 runs)` by Firth’s penalised logistic regression [37, 38], with sequencing depth and study size as the effort covariates. Fits used `brglm2` v1.1.0 `brglmFit` with `type` `=` `"AS_mean"` in R v4.3.3 and the FLIC intercept correction [39], and 95% confidence intervals were computed by penalised profile likelihood with `logistf` v1.26.1.

A genus was compared between host nodes only where none of its species falls in a collapse group spanning more than one genus at either deposited V region. Genera were not compared with each other within a single host node. Amplicon region is close to collinear with host node and was not a model term. The detection contrasts at host-class, host-family and host-genus rank were therefore refitted within a single region instead, 335 of the 2,525 contrasts, and a host-class contrast is read only against that refit (Table S6; Supplementary Methods). Contrasts were corrected for multiple testing by the Benjamini-Hochberg procedure within each pre-specified family of tests, over every pair tested in that family rather than every pair reported. The detection contrasts fall into 181 such families, each being one genus within one host parent taxon, and the undescribed-rate contrasts into 50, each being one host parent taxon.

Three reporting rules were applied. An undescribed rate was reported for a host node only where that node has five or more analysable BioProjects. An undescribed OTU entered the reported detections only where it recurred in at least two independent in-scope BioProjects. A genus was estimated at a host node, and was eligible for a contrast there, only where it was detected in at least two BioProjects of that node, so 3,117 of 9,598 genus-by-node cells carry an estimate. The five-BioProject threshold is a working value and the fit was re-run at each candidate value to show its effect (Tables S7a, S7b). The two-BioProject recurrence requirement was not varied.

One BioProject contributes at most one unit per host node, and where a BioProject sampled several hosts its rows were treated as independent (Supplementary Methods). The two compositional quantities are descriptive and are not compared between hosts. Every between-host statement rests on a fitted contrast.

## RESULTS AND DISCUSSION

### Classification outcome

Of the 349,712 OTUs the family-membership test confirmed, 204,813 entered the analysed frame from 3,344 independent BioProjects of plant and invertebrate hosts (Table S8). Of those, 20,516 were named to species and 127,906 to genus, 51,650 were ambiguous, and 4,741 were ruled out of every described species and are reported as undescribed (Figures 1A, 2A; Table S9). Thirty-seven genera were detected, every genus the family holds. A genus-rank call dominates across the 43 plant and 44 invertebrate host nodes of Figures 1A and 2A (Table S10), with species far less resolved, matching an independent assessment in which genus assignment over these regions is accurate and species assignment hardly possible [18].

**Fig. 1.**
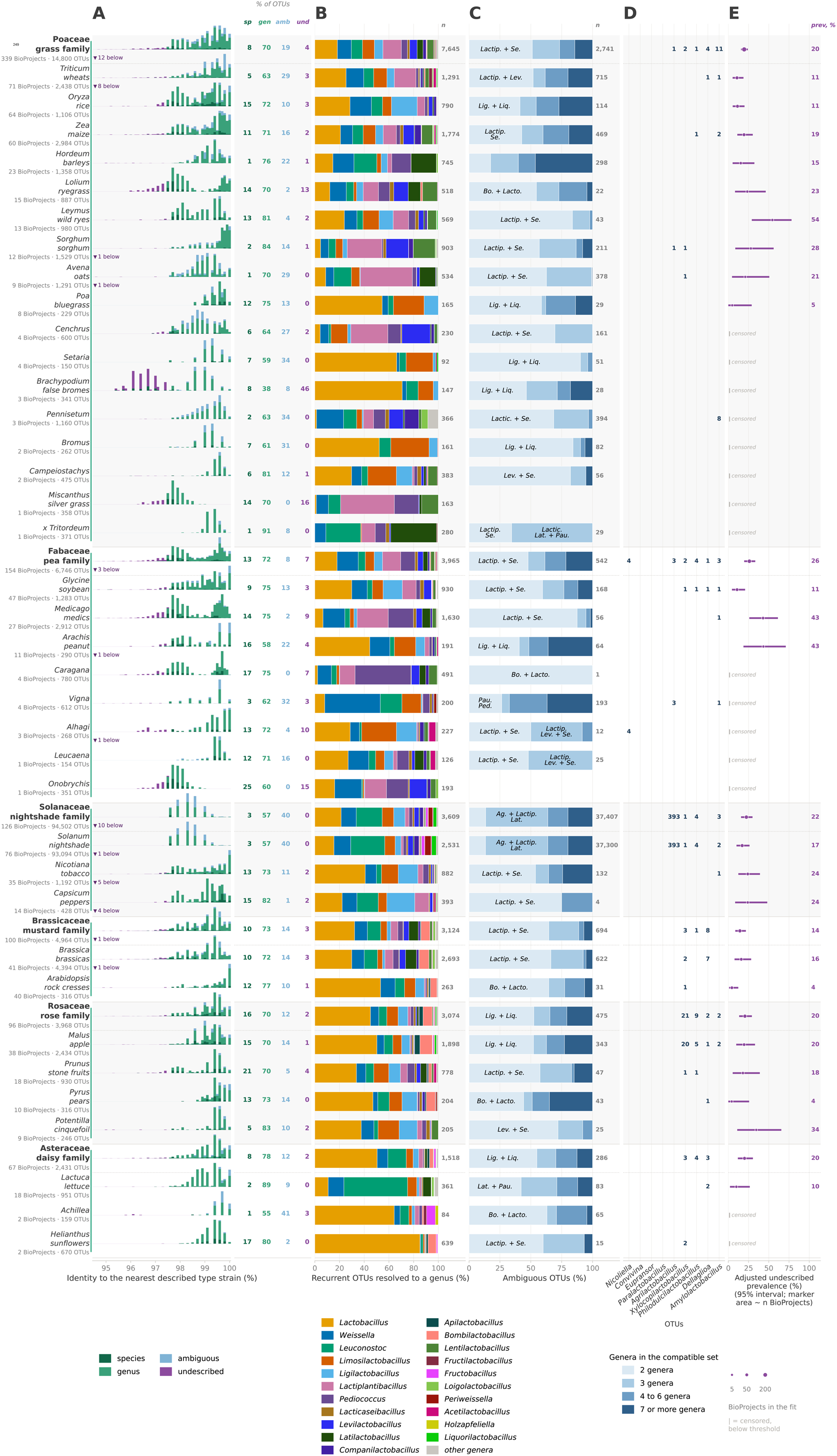
Classification of *Lactobacillaceae* 16S rRNA gene amplicons across plant hosts, by host node. Rows are host families in bold, each followed by its own host genera indented within a shaded band, and a family row contains the rows beneath it. Six families and 37 genera are shown, ordered by the number of independent BioProjects supporting each node, given beneath the node name with the operational taxonomic unit (OTU) count. A node is shown where it carries at least 150 OTUs. An OTU is the seed sequence of a Swarm *d* = 1 cluster. (A) Percent identity of each OTU to its nearest described type strain, in four outcome classes. Each bar is one value of the 0.1-point grid between the 94.5% genus threshold and 100%. Each row is scaled to its own tallest bar. OTUs below 94.5% are counted at the left of the row and are not drawn. Percentages beside the panel give that row’s composition across the four classes. (B) Lineage composition of each row’s named *Lactobacillaceae*, as the share of the OTUs recurring across at least 2 independent in-scope BioProjects and resolved to a genus, with 21 genera carrying a colour and all remaining genera pooled. Species-rank calls are counted under their own genus. **n** beside each bar is the number of recurrent OTUs resolved to a genus. Genera are drawn in one order on both plates, so a genus keeps its colour and its position whichever figure it appears in. A family row and the genus rows beneath it are separated by a dashed rule. (C) Number of genera in the compatible set of each row’s ambiguous OTUs. An ambiguous OTU has a non-empty compatible set spanning more than one genus. Bands are ordinal and **n** beside each bar is the number of ambiguous OTUs. (D) OTU counts for nine rare genera, being those whose share never reaches 3% of a row’s genus-resolved OTUs on either plate, so no segment of them in (B) would be legible. The counts are also deposited per genus and host node in Table S13a. A blank cell is an absence of OTUs. (E) Adjusted undescribed prevalence, the proportion of a node’s BioProjects yielding an undescribed OTU, with 95% penalised profile-likelihood intervals, from Firth penalised logistic regression with the FLIC intercept correction, fitted per BioProject and adjusted for sequencing depth and study size. Marker area scales with the number of BioProjects in the fit. 15 of the 43 nodes fall below the reporting threshold and are marked censored at the axis foot. An OTU is counted once per drawn row and the rows nest, so a total summed down a panel exceeds the number of distinct OTUs behind it. The 2,818 undescribed OTU-row counts drawn here rest on 2,318 distinct undescribed OTUs. Genus names inside the bands are abbreviated as follows: *Agrilactobacillus* Ag., *Amylolactobacillus* Am., *Apilactobacillus* Ap., *Bombilactobacillus* Bo., *Companilactobacillus* Com., *Convivina* Con., *Daquilactobacillus* Da., *Dellaglioa* De., *Eupransor* Eu., *Fructilactobacillus* Fructi., *Fructobacillus* Fructo., *Holzapfeliella* Ho., *Lacticaseibacillus* Lactic., *Lactiplantibacillus* Lactip., *Lactobacillus* Lacto., *Lapidilactobacillus* Lap., *Latilactobacillus* Lat., *Lentilactobacillus* Len., *Leuconostoc* Leu., *Levilactobacillus* Lev., *Ligilactobacillus* Lig., *Limosilactobacillus* Lim., *Liquorilactobacillus* Liq., *Loigolactobacillus* Lo., *Nicoliella* Ni., *Paucilactobacillus* Pau., *Pediococcus* Ped., *Philodulcilactobacillus* Ph., *Schleiferilactobacillus* Sc., *Secundilactobacillus* Se., *Weissella* We. **Alt text:** A five-panel plate of plant hosts, one row per host node: 6 host families each followed by its own indented genera, 43 rows in all, ordered top to bottom by the number of independent BioProjects behind each node. Panel A gives each row a histogram of percent identity to the nearest described type strain, from the 94.5% genus boundary on the left to 100% on the right, bars coloured by classification outcome and each row scaled to its own tallest bar; most rows pile up against the right-hand edge, and OTUs below the boundary are tallied at the left rather than drawn. Panel B is a matrix of the share of each row’s named lineages going to each genus. Panel C is a stacked bar per row of how many genera the compatible set spanned, shaded light to dark. Panel D tabulates OTU counts for 9 rare genera. Panel E plots adjusted undescribed prevalence per row with intervals, marker area scaling with the BioProjects in the fit and 15 rows censored at the axis foot.

**Fig. 2.**
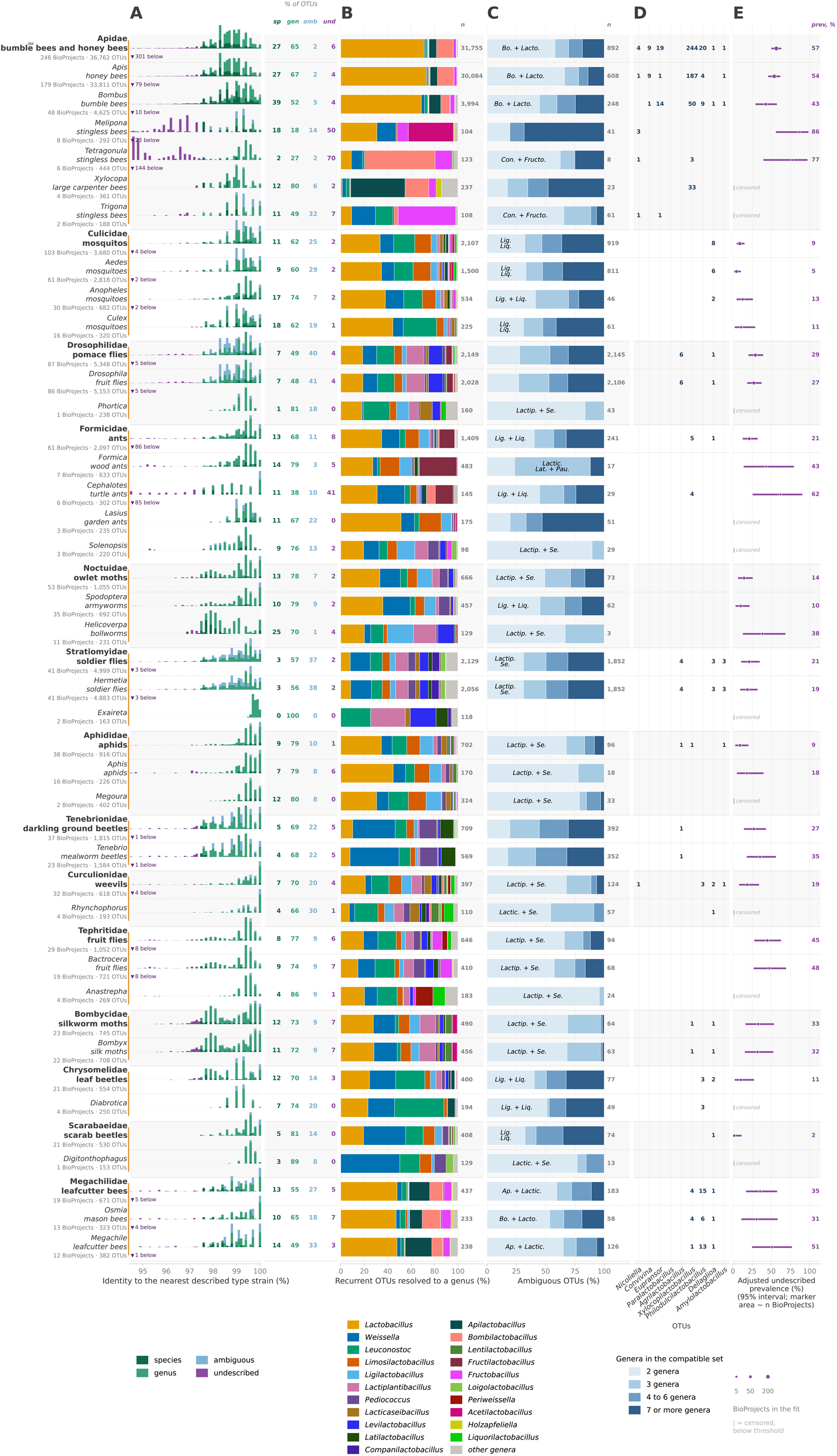
Classification of *Lactobacillaceae* 16S rRNA gene amplicons across invertebrate hosts, by host node. Rows are host families in bold, each followed by its own host genera indented within a shaded band, and a family row contains the rows beneath it. 14 families and 30 genera are shown, ordered by the number of independent BioProjects supporting each node, given beneath the node name with the operational taxonomic unit (OTU) count. A node is shown where it carries at least 150 OTUs. An OTU is the seed sequence of a Swarm *d* = 1 cluster. (A) Percent identity of each OTU to its nearest described type strain, in four outcome classes. Each bar is one value of the 0.1-point grid between the 94.5% genus threshold and 100%. Each row is scaled to its own tallest bar. OTUs below 94.5% are counted at the left of the row and are not drawn. Percentages beside the panel give that row’s composition across the four classes. (B) Lineage composition of each row’s named *Lactobacillaceae*, as the share of the OTUs recurring across at least 2 independent in-scope BioProjects and resolved to a genus, with 21 genera carrying a colour and all remaining genera pooled. Species-rank calls are counted under their own genus. **n** beside each bar is the number of recurrent OTUs resolved to a genus. Genera are drawn in one order on both plates, so a genus keeps its colour and its position whichever figure it appears in. A family row and the genus rows beneath it are separated by a dashed rule. (C) Number of genera in the compatible set of each row’s ambiguous OTUs. An ambiguous OTU has a non-empty compatible set spanning more than one genus. Bands are ordinal and **n** beside each bar is the number of ambiguous OTUs. (D) OTU counts for nine rare genera, being those whose share never reaches 3% of a row’s genus-resolved OTUs on either plate, so no segment of them in (B) would be legible. The counts are also deposited per genus and host node in Table S13a. A blank cell is an absence of OTUs. (E) Adjusted undescribed prevalence, the proportion of a node’s BioProjects yielding an undescribed OTU, with 95% penalised profile-likelihood intervals, from Firth penalised logistic regression with the FLIC intercept correction, fitted per BioProject and adjusted for sequencing depth and study size. Marker area scales with the number of BioProjects in the fit. Eleven of the 44 nodes fall below the reporting threshold and are marked censored at the axis foot. An OTU is counted once per drawn row and the rows nest, so a total summed down a panel exceeds the number of distinct OTUs behind it. The 5,863 undescribed OTU-row counts drawn here rest on 3,261 distinct undescribed OTUs. Genus names inside the bands are abbreviated as follows: *Agrilactobacillus* Ag., *Amylolactobacillus* Am., *Apilactobacillus* Ap., *Bombilactobacillus* Bo., *Companilactobacillus* Com., *Convivina* Con., *Daquilactobacillus* Da., *Eupransor* Eu., *Fructilactobacillus* Fructi., *Fructobacillus* Fructo., *Furfurilactobacillus* Fu., *Holzapfeliella* Ho., *Lacticaseibacillus* Lactic., *Lactiplantibacillus* Lactip., *Lactobacillus* Lacto., *Lapidilactobacillus* Lap., *Latilactobacillus* Lat., *Lentilactobacillus* Len., *Leuconostoc* Leu., *Levilactobacillus* Lev., *Ligilactobacillus* Lig., *Limosilactobacillus* Lim., *Liquorilactobacillus* Liq., *Loigolactobacillus* Lo., *Nicoliella* Ni., *Paralactobacillus* Par., *Paucilactobacillus* Pau., *Pediococcus* Ped., *Philodulcilactobacillus* Ph., *Schleiferilactobacillus* Sc., *Secundilactobacillus* Se., *Weissella* We. **Alt text:** A five-panel plate of invertebrate hosts, one row per host node: 14 host families each followed by its own indented genera, 44 rows in all, ordered top to bottom by the number of independent BioProjects behind each node. Panel A gives each row a histogram of percent identity to the nearest described type strain, from the 94.5% genus boundary on the left to 100% on the right, bars coloured by classification outcome and each row scaled to its own tallest bar; most rows pile up against the right-hand edge, and OTUs below the boundary are tallied at the left rather than drawn. Panel B is a matrix of the share of each row’s named lineages going to each genus. Panel C is a stacked bar per row of how many genera the compatible set spanned, shaded light to dark. Panel D tabulates OTU counts for 9 rare genera. Panel E plots adjusted undescribed prevalence per row with intervals, marker area scaling with the BioProjects in the fit and 11 rows censored at the axis foot.

The ambiguous class is overwhelmingly a plant-host phenomenon, and narrow rather than diffuse (Figures 1C, 2C; Table S11). Of the 51,650 ambiguous OTUs, 45,924 occur in plant hosts and 9,753 in invertebrate hosts, 4,027 in both. The ambiguity takes a different shape in each host class, 44.4% of the plant ambiguity standing between exactly three genera and the invertebrate ambiguity being mostly a choice between two genera, at 40.0%. One three-genus set, *Agrilactobacillus*, *Lactiplantibacillus* and *Latilactobacillus*, carries 28.8% of the whole ambiguous class and a third of the plant class on its own, while the commonest invertebrate set reaches 7.2% of that class. Those three are not close relatives, each a fully supported clade with their common ancestor at the base of the family (Figure S1).

A panel of 12,378 control, mock and blank runs tested whether the detections below were reagent background (Table S3b). Nine genera shared no OTU seed sequence with any control, and no rare genus discussed below shares a seed with a control from an analysed BioProject. *Apilactobacillus* and *Periweissella* each shared one seed with a control, out of 3,699 and 6,089 OTUs respectively, so reagent background and plate carryover are excluded as the source of what follows.

### Naming failure in published studies

Six published studies reported a *Lactobacillaceae* detection they could not name (Table S12a), spanning 2009 to 2026, three continents and five host systems, one a culture study and five amplicon surveys. Three report identity evidence contradicting their own printed label, and in the other three the contradiction appears only against a type-strain reference. Husseneder *et al*. [40] cultured three lactic acid bacterial morphotypes from every *Coptotermes formosanus* colony sampled, named none to genus or species, and placed one isolate in the Lactobacillales with no GenBank match above 95% identity. Liu *et al*. [41] surveyed 121 nests and hives of *Tetragonula carbonaria* and *Austroplebeia australis* and reported four “*Lactobacillus* spp.” in the core of each, one year after four species were described from those two hosts [42]. Garza-González *et al*. [43] found the single amplicon sequence variant shared by *Melipona beecheii* and *M. yucatanica* assigned to *Lactobacillus bombicola* by SILVA and by the bee-specific BEExact database at below 97% identity to that species. Curating the reference does not close the gap. BEExact carries 618 placeholder labels for uncultivated bee lineages and reaches species rank for 80 to 90% of short-read variants [44], yet returned an unsupported name.

Reclassified under the exclusion rule, every study whose reads were retrieved returns a call those reads support (Table S12b). Liu’s reads resolve to *Bombilactobacillus*, compatible with only two described species, both named from those hosts the year before and inseparable at V3-V4; Garza-González’s *Melipona* reads resolve to *Acetilactobacillus* and not to *L. bombicola*; and most *Cephalotes* OTUs are undescribed. Husseneder’s isolates predate the archive, and Mills’s runs carry a metagenome taxon this survey does not retrieve. In four of the six the printed name is wrong at the rank it names rather than merely short of one, and no correction needed new sequencing.

### Host range of the rare genera

Six genera known from one or two isolates and one habitat recur across many independent BioProjects (Figures 1B, 1D, 2B, 2D). *Xylocopilactobacillus* was described from Japanese *Xylocopa* carpenter bees and framed as *Xylocopa*-restricted and vertically transmitted [45], a framing restated in 2026 [46]. It was detected here in 163 BioProjects, at a fitted prevalence of 34.0% in Apidae against 3.4% in Formicidae (odds ratio 14.6, 95% CI 5.42 to 54.5, Benjamini-Hochberg adjusted *p* < 0.0001; Tables S13a, S13b). Of its detections 472 resolve to *X. apis* or *X. apicola*, and *Xylocopa* itself carries 33 *Xylocopilactobacillus* OTUs, fifteen resolving to species, in one of its four BioProjects. *Philodulcilactobacillus myokonensis*, from fermented vegetable extracts in Niigata [47, 48], was detected across solitary and wild bees in 45 BioProjects, every detection at species rank, at 1.4% in *Apis* against 45.3% in *Ceratina* (odds ratio 0.016, 95% CI 0.002 to 0.091; Table S13b). *Paralactobacillus selangorensis*, from a Malaysian fermented food ingredient, was recovered at species rank from *Periplaneta americana* guts in six BioProjects, and *Acetilactobacillus jinshanensis*, from a Chinese grain vinegar mash, in 145 BioProjects with every one of its 1,396 detections at species rank [2, 49]. *Eupransor* and *Convivina*, from bumble bee guts and a hornet hindgut [50–52], recur across four *Vespa* BioProjects in Europe and eastern Asia, and *Convivina* reaches the Mexican honey wasp *Brachygastra mellifica* on 21 OTUs. Holley *et al*. [53] report it there independently, alongside a *Bombilactobacillus* they place close to *B. apium*. This survey’s own reads from that host resolve to the same species, on 15 OTUs.

Hammer *et al*. [54] supply a potential mechanism, showing that cellophane bee larval provisions become near-monocultures of lactobacilli, the host engineering a sugar-rich, high-water, actively fermenting microenvironment in which already-widespread generalists dominate rather than new symbionts being acquired. A vegetable ferment, a grain vinegar mash and a bee provision are one environment made three ways, and it selects a fructophilic, acidophilic, sucrose-tolerant organism [8, 47]. A species description names the substrate that yielded a colony on the medium then in use, so the habitat it names is a lower bound on host range. Isolation source and frequency of isolation are two of the five criteria Duar *et al*. [1] assigned this family’s lifestyles from, and the six genera above change both.

### Bee-associated genera in plant hosts

The three genera named from bees are not spread evenly across plant families, and their distribution follows pollination rather than sampling effort (Figures 1B, 1D; Table S13a). Rosaceae and Brassicaceae carry 230 and 230 *Bombilactobacillus* OTUs and 94 and 19 *Apilactobacillus* OTUs, against 41 and 21 for Poaceae. Poaceae is the most heavily sampled of the three at 339 BioProjects to Rosaceae’s 96, and *Xylocopilactobacillus* follows the same pattern at 21, 3 and 2. The fitted rates agree, *Bombilactobacillus* being detected in 17.3% of Rosaceae BioProjects (11.2 to 25.2) against 3.1% of Poaceae ones (1.8 to 5.0), and *Apilactobacillus* in 12.4% (7.3 to 19.1) against 1.8% (0.9 to 3.3), neither pair of intervals overlapping. No fitted contrast between plant families is deposited for these genera, so these are two rates read side by side, not an odds ratio.

The same three genera are detected more often in an invertebrate study than a plant one, at odds ratios of 9.05 for *Xylocopilactobacillus*, 5.05 for *Apilactobacillus* and 4.14 for *Bombilactobacillus*, each in the same direction inside both amplicon regions with every interval excluding one (Tables S6, S13b). *Weissella* gives 1.01 and separates the two host classes in neither region. *Furfurilactobacillus* gives 0.54 in V3-V4 and 2.37 in V4, so its pooled value of 1.07 is two opposed region effects cancelling and supports no statement about host class.

Insect pollination separates those families, and bees carry these bacteria between flowers. Argueta-Guzmán *et al*. [55] showed experimentally that *Osmia lignaria* moves *Apilactobacillus micheneri* from a flower to a previously uninoculated one, where a second bee picks it up. Floral microbiomes carry bee-associated genera at lower abundance than the bees themselves [56]. Anderson *et al*. [57] noticed core honey bee gut lactobacilli in another group’s apple flower data a decade ago. The BioSample sample-type annotation agrees. Of the 378 Rosaceae detections of these three genera, 34.1% are annotated phyllosphere and 21.2% flower, the other 44.7% carrying no annotation, so every annotated detection is a leaf surface or a flower and none a root, a seed or soil. We found no published report of *Bombilactobacillus* from a plant, and 16S surveys of apple and pear floral nectar named no *Lactobacillaceae* genus at all [58]. A genus with no plant record in the literature is therefore present across two crop families here, recovered only because every read was recruited against a type-strain reference first.

### Phenotype and habitat

Three habitat associations are predictable from the type strains, and the fit recovered all three from the amplicon data alone. *Amylolactobacillus*, whose two species are amylolytic and were isolated from starch-rich fermentation [2], reaches 9.1% in Blattodea against 0.3% in Hymenoptera (odds ratio 31.4, 95% CI 7.32 to 182.5; Table S13b), and the American cockroach thrives on a plant-polysaccharide-rich diet whose cultivable gut symbionts degrade starch [59]. *Dellaglioa*, two of whose three species were described from packaged chilled beef, reaches 17.8% in Calliphoridae against 1.4% in Culicidae (odds ratio 15.2; Table S13b), and blowfly larvae develop in decomposing meat. The bee fructophiles behave as the cultivation record predicts, *Apilactobacillus* reaching 67.9% in Apidae against 10.3% in Formicidae (odds ratio 18.4) and *Bombilactobacillus* 66.5% against 8.9% (odds ratio 20.4). Both rise inside the V4 region alone, to odds ratios of 36.6 and 29.6 (Table S6), so primer choice is not a confounder. Of 2,525 tested contrasts, 152 survive Benjamini-Hochberg correction and fall on 13 genera, eleven neither *Lactobacillus* nor *Weissella*, the two most prevalent. No isolation source or phenotype enters the model, so all three associations were recovered without being sought.

### *Fructilactobacillus* in honeydew-feeding ants

Undescribed OTUs nearest *Fructilactobacillus* give the deepest undescribed signals here, and it is the one genus in which the same species is recorded from insects and from foods. Zheng *et al*. [60] sequenced the digestive tracts of three *Formica* species and *Lasius niger*, all aphid-honeydew feeders, and found lactic acid bacteria concentrated in the infrabuccal pockets and crops. They isolated *Lactobacillus sanfranciscensis* and *L. lindneri*, the first of which catabolises the honeydew sugars sucrose, trehalose, melezitose and raffinose. Both are *Fructilactobacillus* under the 2020 framework, which that paper does not apply. *Formica* carries 633 OTUs here, *Fructilactobacillus* holding 223 of them across six of its seven BioProjects, including 31 called as *F. lindneri*. The genus also reaches the honeydew-tending weaver ant *Oecophylla smaragdina*, in two BioProjects from Malaysia and Cambodia for which this family was not a focus. Three of those OTUs are undescribed, each nearest to *F. myrtifloralis* [61] at 96.6 to 96.8% identity. Our own collection reaches the genus too. Of 714 isolates from wild Australian niches, 91 are *Fructilactobacillus* and the 31 unassignable to a described species all came from the two green tree ant sites [62]. Together these datasets, spanning different countries, materials and methods, place an undescribed *Fructilactobacillus* in green tree ants, and the strains needed to describe it are already in culture.

### Undescribed lineages in stingless bees

*Melipona* and *Tetragonula* hold the two highest undescribed prevalences of the 52 host nodes drawn in Figure 3, at 86.1% (56.0 to 98.4) and 77.4% (39.7 to 97.2), one Neotropical host and one Indo-Australian. Both also carry a long tail of OTUs below the 94.5% genus threshold (Figure 2A), 23 of *Melipona*’s 292 OTUs and 144 of *Tetragonula*’s 444 falling below it against 79 of *Apis*’s 33,811. *Melipona* lacks the corbiculate core [43, 63], which is an effort-adjusted contrast here. *Lactobacillus* detection is 95.1% in *Apis* against 30.2% in *Melipona* (odds ratio 44.8; Table S13b), and 91.2% in *Bombus* (odds ratio 23.8, 95% CI 4.0 to 191). The separation therefore holds against a second corbiculate host, though the *Bombus* interval is nearly fiftyfold wide on 8 *Melipona* BioProjects. *Weissella* shows no detectable separation between the same two hosts (odds ratio 0.71, 95% CI 0.16 to 4.08, adjusted *p* = 0.95), so the difference is a property of *Lactobacillus* and not of how deeply *Melipona* was sequenced. *Acetilactobacillus* is instead its most OTU-rich named genus, at 41 of 292 OTUs, and one of the environmentally acquired *Lactobacillaceae* Cerqueira *et al*. [63] describe in this host. A further 145 of the 292 are undescribed. *Tetragonula* keeps its socially transmitted core, retaining *Bombilactobacillus* on 74 OTUs across four of its six BioProjects while still carrying 309 undescribed OTUs, 171 anchored to *Acetilactobacillus*, so one host can hold both.

**Fig. 3.**
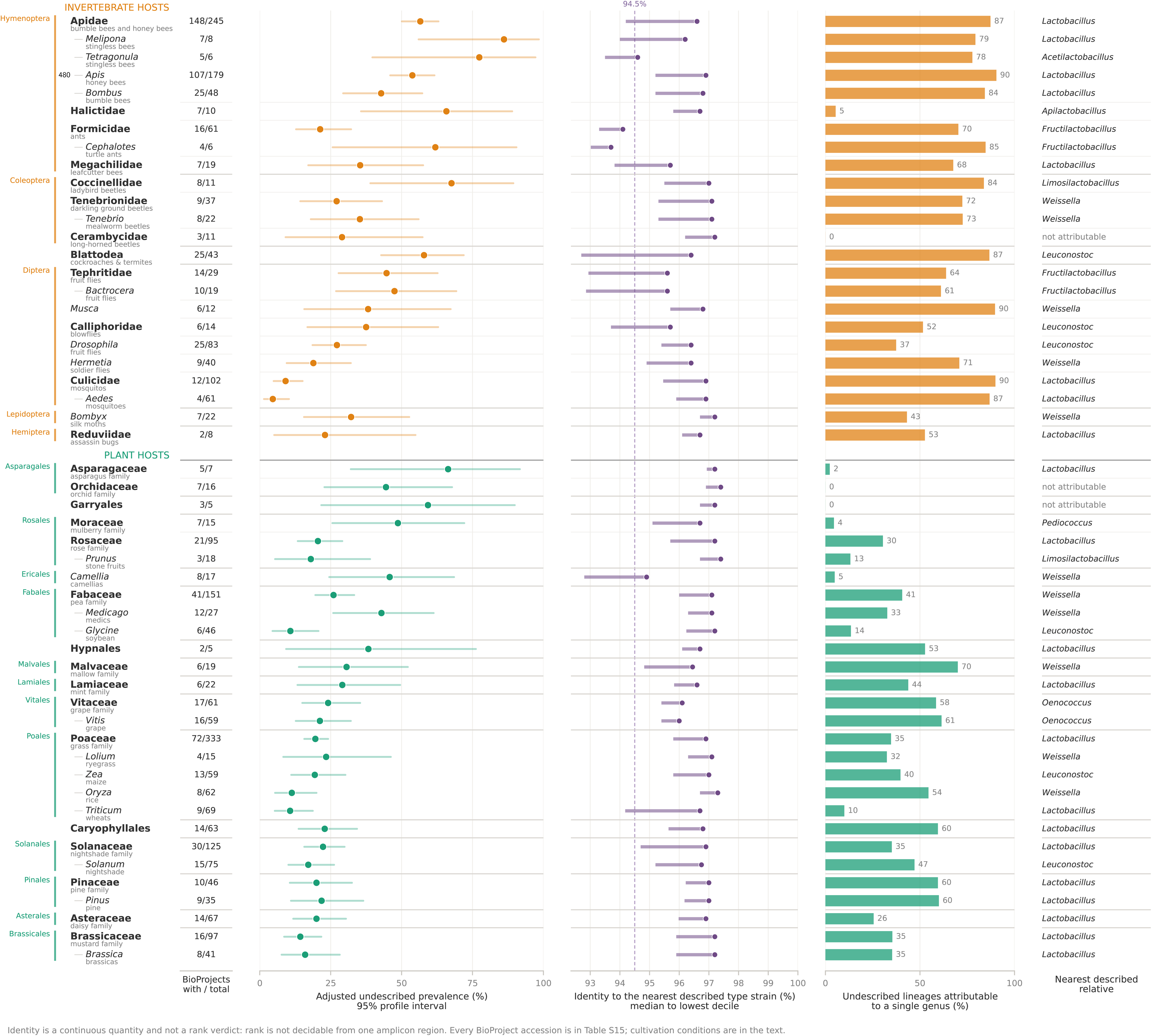
Prevalence, percent identity and genus attribution of undescribed *Lactobacillaceae*, by host taxon. 52 host taxa, 24 invertebrate and 28 plant, each carrying at least 30 distinct undescribed operational taxonomic units (OTUs), ranked by adjusted undescribed prevalence within host class. An OTU is the seed sequence of a Swarm *d* = 1 cluster. Rows are given at the rank each lineage is reported at; where a family and its only sampled genus carry the same observations the finer rank is kept, reducing 58 candidate rows to 52. BioProjects with / total: independent studies of that host in which an undescribed lineage was detected, over studies sampled. Adjusted undescribed prevalence: the proportion of a host’s BioProjects carrying at least one undescribed lineage, adjusted for sequencing depth and run count, with a 95% penalised profile-likelihood interval. It is the quantity panel (E) of Figures 1 and 2 draws. Percent identity to the nearest described type strain: matched positions between each undescribed OTU and its nearest described type strain over the aligned amplicon window, drawn from the median out to the lowest decile. The axis ascends to 100% on the right, as in Figures 1 and 2. The 94.5% mark is the genus-level reference of Yarza *et al*. [31], drawn for orientation and not applied as a classifier. Attributability: the share of a host’s undescribed OTUs whose nearest described type strain resolves to a single genus at that OTU’s own amplicon region. An OTU whose nearest relative sits in a collapse group spanning several genera is left unassigned. Every BioProject accession behind every row is in Table S15, and the within-project control analysis is Table S16. **Alt text:** A table-style figure with 52 rows, one per host taxon, split into an invertebrate block of 24 rows and a plant block of 28 rows and ranked within each block by adjusted undescribed prevalence, highest at the top. Each row carries the host name and then four aligned columns: the number of BioProjects with an undescribed lineage over the number sampled; a dot-and-interval plot of adjusted undescribed prevalence from 0 to 100 per cent; a horizontal range running from each host’s median percent identity to the nearest described type strain out to its lowest decile, on an axis ascending to 100 per cent on the right with a vertical reference line at 94.5 per cent; and a bar of the share of that host’s undescribed OTUs attributable to a single genus. Melipona sits at the top of the prevalence column and Cephalotes furthest left on the identity column, the two extremes being different rows.

Shotgun metagenomics independently reaches the same two genera in the same hosts. Xolalpa-Aroche *et al*. [64] assembled 24 metagenome-assembled genomes from 17 *Melipona beecheii* and *Scaptotrigona mexicana* pot honey samples. Fifteen sat at or below 81% average nucleotide identity to any described species, resolving into four clades consistent with four undescribed species, two nearest *Nicoliella* and two nearest *Acetilactobacillus* (Figure 2B). Their design carries whole genomes from two Mexican bee species and no amplicon data. This study carries 17 stingless bee BioProjects and no genomes. For two of the four clades their genomes already supply the confirmation an amplicon call inherently lacks. Seven undescribed OTUs here are anchored to *Nicoliella*, two of them in *Melipona*, so their *Nicoliella*-adjacent clades and this survey point at the same undescribed lineage.

*Nicoliella* occurs in both host regions (Figure 2D), though only two species are described in it, from Australian stingless bees and from *Lavandula angustifolia* flowers, and no culture study has reported it from the Neotropics [42, 65]. Sarton-Lohéac *et al*. [66], sequencing gut symbionts of six Neotropical stingless bee species, cite it once and did not recover it. It was detected here in 12 BioProjects on 8 OTUs and 26 detections. Its stingless bee records span six genera, *Tetragonula* and five Neotropical ones, on 4 OTUs and 9 detections, which establishes occurrence but not abundance. *Tetragonula carbonaria*, the type host of *Nicoliella spurrieriana*, *Bombilactobacillus folatiphilus* and *B. thymidiniphilus* [42, 67], is both the most productive described source of this family among stingless bees and one of its richest undescribed sources. All three were named in one 2022 study, and the naming is not finished. Halictidae, the sweat bees, yields an undescribed OTU in 65.8% of its ten BioProjects (35.7 to 89.0), on 74 undescribed OTUs (Figure 3).

### Divergence and the limits of the survey

*Cephalotes* carries the lowest median identity to a nearest described type strain of those 52 host nodes, at 93.7% across its undescribed OTUs, with 61.9% undescribed prevalence, 124 undescribed OTUs and *Fructilactobacillus* as the closest genus. Turtle ants hold one of the better-characterised insect gut communities described, its core persisting at least 50 million years, with the family in it at 2%, named only as “*Lactobacillus*” [15, 68, 69]. Formicidae as a whole carries the second lowest median identity at 94.1%, so the signal is not confined to one ant genus. Four of the five lowest median identities are invertebrate nodes, the one exception being tea, *Camellia*, at 94.9% on 61 undescribed OTUs, 58 of them left unassigned at genus. At family rank, Apidae yields an undescribed OTU in 56.6% of its BioProjects against 21.3% for Formicidae (odds ratio 4.81; Table S14), and that ratio holds between 4.74 and 4.87 at every reporting threshold from 5 to 30 analysable BioProjects (Tables S7a, S7b). The most divergent *Lactobacillaceae* found here therefore sit in a community sequenced repeatedly without anyone naming them, making *Cephalotes* the most attractive target for formal description.

Of the 1,754 undescribed OTUs left unassigned at genus, 827 came from a V region with no deposited collapse groups and 927 from a neighbourhood spanning more than one genus, so the failure to assign is a property of the primer and the marker rather than of the lineage (Supplementary Methods). Sampling is also uneven, so no absence is claimed. A run whose deposited metadata names no host at all is out of scope (Supplementary Methods). Figure 3 draws only the host nodes carrying at least thirty undescribed OTUs. Every node clearing the reporting threshold is in Table S15. Most host pairs sampled inside one BioProject agree in their undescribed percentage (Table S16), so the prospecting rates are read as a description and no host-to-host comparison is drawn.

### Cultivation requirements

Forty-eight described species have a supplemented routine or isolation medium in their primary descriptions, 39 of the 52 species in the nine surveyed genera and nine species of *Lactobacillus* from bees. Four of the eight genera represented carry a host contrast that survives correction here (Tables S1, S13b). D-fructose is the commonest supplement, in the routine medium of 31 of the 48, supplied as an electron acceptor to strains whose *adhE* gene is incomplete or absent [7, 70]. L-cysteine is in the medium of 16, eleven of which also carry fructose, every one from a bee gut, a honey stomach, bee bread, a whole bee, an ant or a cockroach, and none from a plant source [71, 72]. The most fastidious were recovered on undefined plant juices, *Bombilactobacillus folatiphilus* reaching no appreciable density in MRS until 20% apple juice was added and *Fructilactobacillus fructivorans* none until tomato juice or fructose [42, 73], and both *Xylocopilactobacillus* species lack the genes for NAD biosynthesis [45]. Culturing undescribed *Lactobacillaceae* from these hosts will therefore require supplementation.

## Conclusion

Applying a type-anchored exclusion rule to 3,344 public BioProjects returns three results. Datasets whose authors could name their *Lactobacillaceae* only as “*Lactobacillus*”, or not at all, are reclassified to the rank their own reads support. The habitat a species was described from predicts its phenotype and not its host range. The deepest lineages are in the hosts with the fewest cultured representatives. *Melipona* and *Tetragonula* carry undescribed *Lactobacillaceae* in most studies that sample them, and *Cephalotes* carries the most divergent lineages. Those hosts are also the hardest to culture, and for forty-eight described species from them the primary description specifies at least one supplement, so bioprospecting in this family now has a shortlist of target hosts and an initial medium for each (Tables S15, S1).

A formal description reports the medium that yielded its type strain, and similar recipes generally culture similar species. Olofsson *et al*. [71] described four species of the honey bee *Lactobacillus* clade on one medium, MRS with cysteine and fructose. The honey wasp study above [53] cultured its gut bacteria on blood agar and on unsupplemented MRS, isolating *Fructilactobacillus vespulae*, whose own description likewise specifies none. The study did not culture the *Lactobacillus* its authors place in that clade, which this survey detects but assigns to no genus.

*Cephalotes*, *Melipona* and *Tetragonula* should be plated on supplemented media and the isolates carried through to formal description, providing the genomes an amplicon call inherently lacks and beginning to close the reference gap that has made these organisms invisible in studies that have already sequenced them.

## Supporting information

Supplementary File Summaries

## SUPPLEMENTARY MATERIAL

Supplementary material accompanies this preprint.

## ACKNOWLEDGEMENTS

We thank Dr Fabien Voisin, Senior HPC Application Specialist, Division of Research and Innovation, Adelaide University, for consulting during development of the retrieval and recruitment code. We thank the research groups whose sequencing data are deposited in the Sequence Read Archive, on which this study rests.

## AUTHOR CONTRIBUTIONS

**Scott A. Oliphant** (Conceptualization, Data curation, Formal analysis, Investigation, Methodology, Resources, Software, Validation, Visualization, Writing - original draft, Writing - review and editing), **Jennifer M. Gardner** (Funding acquisition, Supervision, Writing - review and editing), **Vladimir Jiranek** (Funding acquisition, Supervision, Writing - review and editing), **Krista M. Sumby** (Conceptualization, Funding acquisition, Methodology, Project administration, Resources, Supervision, Validation, Writing - review and editing).

## DECLARATION OF GENERATIVE AI AND AI-ASSISTED TECHNOLOGIES

During the preparation of this work, the authors used Claude Code (Anthropic, Claude Opus 5 model) to assist in writing the R, Python and shell code for the recruitment and classification pipelines, statistical analysis modules, and figure and table generation scripts. All reported claims and quantities are data derived and not from the model. Full code is deposited (See Data Availability). During code development, the authors reviewed and edited the scripts as needed and take full responsibility for the content of the published article.

## CONFLICTS OF INTEREST

The authors declare that they have no known competing financial interests or personal relationships that could have appeared to influence the work reported in this paper.

## DATA AVAILABILITY

Every sequence analysed is public in the NCBI Sequence Read Archive. The run and BioProject accession of each of the 948,386 runs retrieved, with its per-run recruitment outcome, and the BioProject composition of the analysed sampling frame, are deposited at https://doi.org/10.5281/zenodo.21899558, which also holds the classified units, the amplicon region and nearest-type-strain identity of each, the host resolution, the analysed frame and every fitted model. The sequence of every one of the 349,712 classified OTUs is deposited there as a single FASTA, each carrying its own outcome, the genera its reads are compatible with, the species where one was called, and its identity to the nearest described type strain, so an OTU reported here can be searched, tested against a species described later, and used to design primers. Analysis code is at https://github.com/solipha/lactobacillaceae-plant-invertebrate-amplicon-survey. The type-strain reference is LactoTypeDB v1.0.0, archived at https://doi.org/10.5281/zenodo.20673517 with its build pipeline at https://github.com/solipha/lactotypedb [4].

## Notes

https://doi.org/10.5281/zenodo.21899558

https://github.com/solipha/lactobacillaceae-plant-invertebrate-amplicon-survey

https://doi.org/10.5281/zenodo.20673517

https://github.com/solipha/lactotypedb

### Competing Interest Statement

The authors have declared no competing interest.

## REFERENCES

1. Duar RM, Lin XB, Zheng J et al. Lifestyles in transition: evolution and natural history of the genus Lactobacillus. FEMS Microbiol Rev 2017;41:S27–S48. 10.1093/femsre/fux030

2. Zheng J, Wittouck S, Salvetti E et al. A taxonomic note on the genus Lactobacillus: description of 23 novel genera, emended description of the genus Lactobacillus Beijerinck 1901, and union of Lactobacillaceae and Leuconostocaceae. Int J Syst Evol Microbiol 2020;70:2782–858. 10.1099/ijsem.0.004107

3. Parte AC, Sardà Carbasse J, Meier-Kolthoff JP et al. List of Prokaryotic names with Standing in Nomenclature (LPSN) moves to the DSMZ. Int J Syst Evol Microbiol 2020;70:5607–12. 10.1099/ijsem.0.004332

4. Oliphant SA, Gardner JM, Jiranek V et al. LactoTypeDB v1.0.0: a type-strain 16S rRNA reference database for Lactobacillaceae and its reproducible build pipeline. Zenodo, 2026. 10.5281/zenodo.20673517

5. Kwong WK, Moran NA. Gut microbial communities of social bees. Nat Rev Microbiol 2016;14:374–84. 10.1038/nrmicro.2016.43

6. Endo A, Salminen S. Honeybees and beehives are rich sources for fructophilic lactic acid bacteria. Syst Appl Microbiol 2013;36:444–8. 10.1016/j.syapm.2013.06.002

7. Endo A, Maeno S, Tanizawa Y et al. Fructophilic lactic acid bacteria, a unique group of fructose-fermenting microbes. Appl Environ Microbiol 2018;84:e01290–18. 10.1128/AEM.01290-18

8. McFrederick QS, Vuong HQ, Rothman JA. Lactobacillus micheneri sp. nov., Lactobacillus timberlakei sp. nov. and Lactobacillus quenuiae sp. nov., lactic acid bacteria isolated from wild bees and flowers. Int J Syst Evol Microbiol 2018;68:1879–84. 10.1099/ijsem.0.002758

9. McFrederick QS, Cannone JJ, Gutell RR et al. Specificity between lactobacilli and hymenopteran hosts is the exception rather than the rule. Appl Environ Microbiol 2013;79:1803–12. 10.1128/aem.03681-12

10. Walter J, O’Toole PW. Microbe Profile: the Lactobacillaceae. Microbiology (Reading) 2023;169:001414. 10.1099/mic.0.001414

11. Kang JP, Huo Y, Hoang VA et al. Bombilactobacillus apium sp. nov., isolated from the gut of honeybee (Apis cerana). Arch Microbiol 2021;203:2193–8. 10.1007/s00203-021-02249-y

12. Li TT, Gu CT. Apilactobacillus zhangqiuensis sp. nov. and Apilactobacillus xinyiensis sp. nov., isolated from the gut of honeybee (Apis mellifera). Int J Syst Evol Microbiol 2022;72:005402. 10.1099/ijsem.0.005402

13. Tsai LJ, Chen PC, Lan WT et al. Apilactobacillus intestinapis sp. nov., isolated from the honeybee. Int J Syst Evol Microbiol 2025;75:006971. 10.1099/ijsem.0.006971

14. Kanaya R, Maeno S, Hisatomi A et al. Apilactobacillus pseudapinorum sp. nov., isolated from polyfloral honey. Int J Syst Evol Microbiol 2026;76:007027. 10.1099/ijsem.0.007027

15. Ramalho MO, Moreau CS. Untangling the complex interactions between turtle ants and their microbial partners. Anim Microbiome 2023;5:1. 10.1186/s42523-022-00223-7

16. Mills TJT, Nelson TM, Pearson LA et al. Hive transplantation has minimal impact on the core gut microbiome of the Australian stingless bee, Tetragonula carbonaria. Microb Ecol 2023;86:2086–96. 10.1007/s00248-023-02222-w

17. Tanes C, Tu V, Daniel S et al. Unassigning bacterial species for microbiome studies. mSystems 2024;9:e00515–24. 10.1128/msystems.00515-24

18. Parente E, Zotta T, Giavalisco M et al. Metataxonomic insights in the distribution of Lactobacillaceae in foods and food environments. Int J Food Microbiol 2023;391-393:110124. 10.1016/j.ijfoodmicro.2023.110124

19. Shen W, Ren H. TaxonKit: a practical and efficient NCBI taxonomy toolkit. J Genet Genomics 2021;48:844–50. 10.1016/j.jgg.2021.03.006

20. Leinonen R, Sugawara H, Shumway M. The Sequence Read Archive. Nucleic Acids Res 2011;39:D19–D21. 10.1093/nar/gkq1019

21. Katz K, Shutov O, Lapoint R et al. The Sequence Read Archive: a decade more of explosive growth. Nucleic Acids Res 2022;50:D387–D390. 10.1093/nar/gkab1053

22. Martin M. Cutadapt removes adapter sequences from high-throughput sequencing reads. EMBnet J 2011;17:10–2. 10.14806/ej.17.1.200

23. Rognes T, Flouri T, Nichols B et al. VSEARCH: a versatile open source tool for metagenomics. PeerJ 2016;4:e2584. 10.7717/peerj.2584

24. Edgar RC, Flyvbjerg H. Error filtering, pair assembly and error correction for next-generation sequencing reads. Bioinformatics 2015;31:3476–82. 10.1093/bioinformatics/btv

25. Edgar RC. UPARSE: highly accurate OTU sequences from microbial amplicon reads. Nat Methods 2013;10:996–8. 10.1038/nmeth.2604

26. Mahé F, Rognes T, Quince C et al. Swarm: robust and fast clustering method for amplicon-based studies. PeerJ 2014;2:e593. 10.7717/peerj.593

27. Mahé F, Rognes T, Quince C et al. Swarm v2: highly-scalable and high-resolution amplicon clustering. PeerJ 2015;3:e1420. 10.7717/peerj.1420

28. Mahé F, Czech L, Stamatakis A et al. Swarm v3: towards tera-scale amplicon clustering. Bioinformatics 2022;38:267–9. 10.1093/bioinformatics/btab493

29. Edgar RC, Haas BJ, Clemente JC et al. UCHIME improves sensitivity and speed of chimera detection. Bioinformatics 2011;27:2194–200. 10.1093/bioinformatics/btr381

30. Edgar RC. Search and clustering orders of magnitude faster than BLAST. Bioinformatics 2010;26:2460–1. 10.1093/bioinformatics/btq461

31. Yarza P, Yilmaz P, Pruesse E et al. Uniting the classification of cultured and uncultured bacteria and archaea using 16S rRNA gene sequences. Nat Rev Microbiol 2014;12:635–45. 10.1038/nrmicro3330

32. Brosius J, Palmer ML, Kennedy PJ et al. Complete nucleotide sequence of a 16S ribosomal RNA gene from Escherichia coli. Proc Natl Acad Sci USA 1978;75:4801–5. 10.1073/pnas.75.10.4801

33. Nilsen T, Snipen LG, Angell IL et al. Swarm and UNOISE outperform DADA2 and Deblur for denoising high-diversity marine seafloor samples. ISME Commun 2024;4:ycae071. 10.1093/ismeco/ycae071

34. Callahan BJ, McMurdie PJ, Rosen MJ et al. DADA2: high-resolution sample inference from Illumina amplicon data. Nat Methods 2016;13:581–3. 10.1038/nmeth.3869

35. Wang Q, Garrity GM, Tiedje JM et al. Naïve Bayesian classifier for rapid assignment of rRNA sequences into the new bacterial taxonomy. Appl Environ Microbiol 2007;73:5261–7. 10.1128/aem.00062-07

36. Kim M, Oh HS, Park SC et al. Towards a taxonomic coherence between average nucleotide identity and 16S rRNA gene sequence similarity for species demarcation of prokaryotes. Int J Syst Evol Microbiol 2014;64:346–51. 10.1099/ijs.0.059774-0

37. Firth D. Bias reduction of maximum likelihood estimates. Biometrika 1993;80:27–38. 10.1093/biomet/80.1.27

38. Heinze G, Schemper M. A solution to the problem of separation in logistic regression. Stat Med 2002;21:2409–19. 10.1002/sim.1047

39. Puhr R, Heinze G, Nold M et al. Firth’s logistic regression with rare events: accurate effect estimates and predictions?. Stat Med 2017;36:2302–17. 10.1002/sim.7273

40. Husseneder C, Berestecky JM, Grace JK. Changes in composition of culturable bacteria community in the gut of the Formosan subterranean termite depending on rearing conditions of the host. Ann Entomol Soc Am 2009;102:498–507. 10.1603/008.102.0321

41. Liu H, Hall MA, Brettell LE et al. Microbial diversity in stingless bee gut is linked to host wing size and influenced by the environment. J Invertebr Pathol 2023;198:107909. 10.1016/j.jip.2023.107909

42. Oliphant SA, Watson-Haigh NS, Sumby KM et al. Apilactobacillus apisilvae sp. nov., Nicolia spurrieriana gen. nov. sp. nov., Bombilactobacillus folatiphilus sp. nov. and Bombilactobacillus thymidiniphilus sp. nov., four new lactic acid bacterial isolates from stingless bees Tetragonula carbonaria and Austroplebeia australis. Int J Syst Evol Microbiol 2022;72:005588. 10.1099/ijsem.0.005588

43. Garza-González DA, Quezada-Euán JJG, Medina-Medina LA et al. Comparative analysis of the gut microbiota of the sympatric stingless bee species Melipona beecheii and Melipona yucatanica. Braz J Microbiol 2026;57:85. 10.1007/s42770-026-01905-z

44. Daisley BA, Reid G. BEExact: a metataxonomic database tool for high-resolution inference of bee-associated microbial communities. mSystems 2021;6:e00082–21. 10.1128/msystems.00082-21

45. Kawasaki S, Ozawa K, Mori T et al. Symbiosis of carpenter bees with uncharacterized lactic acid bacteria showing NAD auxotrophy. Microbiol Spectr 2023;11:e00782–23. 10.1128/spectrum.00782-23

46. Silva Cerqueira AE, Holley JaC, Hatcher SC et al. Neffella xylocopae gen. nov., sp. nov., a novel host-specific gut symbiont of Xylocopa carpenter bees in the family Orbaceae. Int J Syst Evol Microbiol 2026;76:007119. 10.1099/ijsem.0.007119

47. Kouya T, Ishiyama Y, Ohashi S et al. Philodulcilactobacillus myokoensis gen. nov., sp. nov., a fructophilic, acidophilic, and agar-phobic lactic acid bacterium isolated from fermented vegetable extracts. PLoS One 2023;18:e0286677. 10.1371/journal.pone.0286677

48. Oren A, Göker M. Validation List no. 214. Valid publication of new names and new combinations effectively published outside the IJSEM. Int J Syst Evol Microbiol 2023;73:006080. 10.1099/ijsem.0.006080

49. Leisner JJ, Vancanneyt M, Goris J et al. Description of Paralactobacillus selangorensis gen. nov., sp. nov., a new lactic acid bacterium isolated from chili bo, a Malaysian food ingredient. Int J Syst Evol Microbiol 2000;50:19–24. 10.1099/00207713-50-1-19

50. Praet J, Meeus I, Cnockaert M et al. Novel lactic acid bacteria isolated from the bumble bee gut: Convivina intestini gen. nov., sp. nov., Lactobacillus bombicola sp. nov., and Weissella bombi sp. nov. Antonie van Leeuwenhoek 2015;107:1337–49. 10.1007/s10482-015-0429-z

51. Hettiarachchi A, Cnockaert M, Joossens M et al. Convivina is a specialised core gut symbiont of the invasive hornet Vespa velutina. Insect Mol Biol 2023;32:510–27. 10.1111/imb.12847

52. Botero J, Peeters C, De Canck E et al. Eupransor demetentiae gen. nov., sp. nov., a novel fructophilic lactic acid bacterium from bumble bees. Int J Syst Evol Microbiol 2024;74:006409. 10.1099/ijsem.0.006409

53. Holley JC, Martin AN, Pham AT, Schlauch J, Moran NA. Honey wasps differ from other wasps in possessing large gut communities dominated by host-restricted bacteria. mBio 2025;16:e02608–24. 10.1128/mbio.02608-24

54. Hammer TJ, Kueneman J, Argueta-Guzmán M et al. Bee breweries: the unusually fermentative, lactobacilli-dominated brood cell microbiomes of cellophane bees. Front Microbiol 2023;14:1114849. 10.3389/fmicb.2023.1114849

55. Argueta-Guzmán M, Spasojevic MJ, McFrederick QS. Solitary bees acquire and deposit bacteria via flowers: testing the environmental transmission hypothesis using Osmia lignaria, Phacelia tanacetifolia, and Apilactobacillus micheneri. Ecol Evol 2025;15:e71138. 10.1002/ece3.71138

56. Keller A, McFrederick QS, Dharampal P et al. (More than) Hitchhikers through the network: the shared microbiome of bees and flowers. Curr Opin Insect Sci 2021;44:8–15. 10.1016/j.cois.2020.09.007

57. Anderson KE, Sheehan TH, Mott BM et al. Microbial ecology of the hive and pollination landscape: bacterial associates from floral nectar, the alimentary tract and stored food of honey bees (Apis mellifera). PLoS One 2013;8:e83125. 10.1371/journal.pone.0083125

58. Smessaert J, Van Geel M, Verreth C et al. Temporal and spatial variation in bacterial communities of ‘Jonagold’ apple (Malus x domestica Borkh.) and ‘Conference’ pear (Pyrus communis L.) floral nectar. Microbiologyopen 2019;8:e918. 10.1002/mbo3.918

59. Vera-Ponce de León A, Jahnes BC, Duan J et al. Cultivable, host-specific Bacteroidetes symbionts exhibit diverse polysaccharolytic strategies. Appl Environ Microbiol 2020;86:e00091–20. 10.1128/aem.00091-20

60. Zheng Z, Zhao M, Zhang Z et al. Lactic acid bacteria are prevalent in the infrabuccal pockets and crops of ants that prefer aphid honeydew. Front Microbiol 2022;12:785016. 10.3389/fmicb.2021.785016

61. Oliphant SA, Watson-Haigh NS, Sumby KM et al. Fructilactobacillus cliffordii sp. nov., Fructilactobacillus hinvesii sp. nov., Fructilactobacillus myrtifloralis sp. nov., Fructilactobacillus carniphilus sp. nov. and Fructobacillus americanaquae sp. nov., five novel lactic acid bacteria isolated from insects or flowers of Kangaroo Island, South Australia. Int J Syst Evol Microbiol 2023;73:005730. 10.1099/ijsem.0.005730

62. Oliphant SA, Sumby KM, Gardner JM et al. Wild Australian niches as a source of novel lactic acid bacteria and malolactic fermentation candidates. Int J Food Microbiol 2026;460:111992. 10.1016/j.ijfoodmicro.2026.111992

63. Cerqueira AES, Lima HS, Silva LCF et al. Melipona stingless bees and honey microbiota reveal the diversity, composition, and modes of symbionts transmission. FEMS Microbiol Ecol 2024;100:fiae063. 10.1093/femsec/fiae063

64. Xolalpa-Aroche A, Contreras-Peruyero H, Delgado-Suárez EJ et al. Genome-resolved metagenomics reveals a phylogenetically cohesive Acetilactobacillus-like species complex dominating stingless bee pot honey. ISME Commun 2026;6:ycag063. 10.1093/ismeco/ycag063

65. Alcántara C, Peirotén Á, Ramón-Nuñez LA et al. Nicoliella lavandulae sp. nov., a novel fructophilic Nicoliella species isolated from flowers of Lavandula angustifolia. Int J Syst Evol Microbiol 2024;74:006497. 10.1099/ijsem.0.006497

66. Sarton-Lohéac G, Nunes da Silva CG, Mazel F et al. Deep divergence and genomic diversification of gut symbionts of Neotropical stingless bees. mBio 2023;14:e03538–22. 10.1128/mbio.03538-22

67. Deshmukh UB, Oren A. Proposal of Christiangramia gen. nov., Neomelitea gen. nov. and Nicoliella gen. nov. as replacement names for the illegitimate prokaryotic generic names Gramella Nedashkovskaya et al. 2005, Melitea Urios et al. 2008 and Nicolia Oliphant et al. 2022, respectively. Int J Syst Evol Microbiol 2023;73:005806. 10.1099/ijsem.0.005806

68. Béchade B, Cabuslay CS, Hu Y et al. Physiological and evolutionary contexts of a new symbiotic species from the nitrogen-recycling gut community of turtle ants. ISME J 2023;17:1751–64. 10.1038/s41396-023-01490-1

69. Flynn PJ, D’Amelio CL, Sanders JG et al. Localization of bacterial communities within gut compartments across Cephalotes turtle ants. Appl Environ Microbiol 2021;87:e02803–20. 10.1128/aem.02803-20

70. Endo A, Okada S. Reclassification of the genus Leuconostoc and proposals of Fructobacillus fructosus gen. nov., comb. nov., Fructobacillus durionis comb. nov., Fructobacillus ficulneus comb. nov. and Fructobacillus pseudoficulneus comb. nov. Int J Syst Evol Microbiol 2008;58:2195–205. 10.1099/ijs.0.65609-0

71. Olofsson TC, Alsterfjord M, Nilson B et al. Lactobacillus apinorum sp. nov., Lactobacillus mellifer sp. nov., Lactobacillus mellis sp. nov., Lactobacillus melliventris sp. nov., Lactobacillus kimbladii sp. nov., Lactobacillus helsingborgensis sp. nov. and Lactobacillus kullabergensis sp. nov., isolated from the honey stomach of the honeybee Apis mellifera. Int J Syst Evol Microbiol 2014;64:3109–19. 10.1099/ijs.0.059600-0

72. Wang C, Huang Y, Li L et al. Lactobacillus panisapium sp. nov., from honeybee Apis cerana bee bread. Int J Syst Evol Microbiol 2018;68:703–8. 10.1099/ijsem.0.002538

73. Charlton DB, Nelson ME, Werkman CH. Physiology of Lactobacillus fructivorans sp. nov., isolated from spoiled salad dressing. Iowa State Coll J Sci 1934;9:1–11.

74. Schober I, Koblitz J, Sardà Carbasse J, et al. BacDive in 2025: the core database for prokaryotic strain data. Nucleic Acids Res 2025;53:D748–D756. 10.1093/nar/gkae959

75. Koblitz J, Halama P, Spring S et al. MediaDive: the expert-curated cultivation media database. Nucleic Acids Res 2023;51:D1531–D1538. 10.1093/nar/gkac803

76. Lee MD. GToTree: a user-friendly workflow for phylogenomics. Bioinformatics 2019;35:4162–4. 10.1093/bioinformatics/btz188

77. Darriba D, Posada D, Kozlov AM et al. ModelTest-NG: a new and scalable tool for the selection of DNA and protein evolutionary models. Mol Biol Evol 2020;37:291–4. 10.1093/molbev/msz189

78. Kozlov AM, Darriba D, Flouri T et al. RAxML-NG: a fast, scalable and user-friendly tool for maximum likelihood phylogenetic inference. Bioinformatics 2019;35:4453–5. 10.1093/bioinformatics/btz305

79. Camacho C, Coulouris G, Avagyan V, et al. BLAST+: architecture and applications. BMC Bioinformatics 2009;10:421. 10.1186/1471-2105-10-421

80. Klindworth A, Pruesse E, Schweer T et al. Evaluation of general 16S ribosomal RNA gene PCR primers for classical and next-generation sequencing-based diversity studies. Nucleic Acids Res 2013;41:e1. 10.1093/nar/gks808

81. Sun DL, Jiang X, Wu QL et al. Intragenomic heterogeneity of 16S rRNA genes causes overestimation of prokaryotic diversity. Appl Environ Microbiol 2013;79:5962–9. 10.1128/AEM.01282-13

82. Kozich JJ, Westcott SL, Baxter NT et al. Development of a dual-index sequencing strategy and curation pipeline for analyzing amplicon sequence data on the MiSeq Illumina sequencing platform. Appl Environ Microbiol 2013;79:5112–20. 10.1128/AEM.01043-13

83. Seemann T. barrnap v1.10.5: BAsic Rapid Ribosomal RNA Predictor. https://github.com/tseemann/barrna (9 August 2026, date last accessed).

84. Chun J, Oren A, Ventosa A et al. Proposed minimal standards for the use of genome data for the taxonomy of prokaryotes. Int J Syst Evol Microbiol 2018;68:461–6. 10.1099/ijsem.0.002516

85. Hackmann TJ. Setting new boundaries of 16S rRNA gene identity for prokaryotic taxonomy. Int J Syst Evol Microbiol 2025;75:006747. 10.1099/ijsem.0.006747

86. Parada AE, Needham DM, Fuhrman JA. Every base matters: assessing small subunit rRNA primers for marine microbiomes with mock communities, time series and global field samples. Environ Microbiol 2016;18:1403–14. 10.1111/1462-2920.13023

87. Costea PI, Zeller G, Sunagawa S et al. Towards standards for human fecal sample processing in metagenomic studies. Nat Biotechnol 2017;35:1069–76. 10.1038/nbt.3960

88. Sinha R, Abu-Ali G, Vogtmann E et al. Assessment of variation in microbial community amplicon sequencing by the Microbiome Quality Control (MBQC) project consortium. Nat Biotechnol 2017;35:1077–86. 10.1038/nbt.3981

89. Stoddard SF, Smith BJ, Hein R et al. rrnDB: improved tools for interpreting rRNA gene abundance in bacteria and archaea and a new foundation for future development. Nucleic Acids Res 2015;43:D593–D598. 10.1093/nar/gku1201

90. Louca S, Doebeli M, Parfrey LW. Correcting for 16S rRNA gene copy numbers in microbiome surveys remains an unsolved problem. Microbiome 2018;6:41. 10.1186/s40168-018-0420-9

