## Supplementary Figures and Methods for "Stingless bees, turtle ants and tea plants are rich sources of undescribed *Lactobacillaceae*"

Running title: Undescribed *Lactobacillaceae* by host

Scott A. Oliphant[a,*], Jennifer M. Gardner[a], Vladimir Jiranek[a,b], Krista M. Sumby[a]

[a]School of Agriculture, Food and Wine, Adelaide University, Waite Campus, Urrbrae, South Australia, Australia

[b]School of Biological Sciences, University of Southampton, Life Sciences B85, Southampton, SO17 1BJ, UK

### Supplementary Methods

The main text states every stage of the method with its tool, threshold and citation, and stands on its own. This file carries the parameters, the exceptions to those rules, and the validation measurements behind statements made in the main text. Sections follow the Materials and Methods in order. Tool versions and non-default parameters are in Table S4, and the analysis code with its per-step documentation is in the software deposit. References are numbered in the main list, except the primary species descriptions behind Table S1, which are listed at the end of this file.

#### Cultivation and media record

The species set was fixed before any description was read.

- Tier one is every species the List of Prokaryotic names with Standing in Nomenclature places in *Apilactobacillus*, *Bombilactobacillus*, *Convivina*, *Eupransor*, *Fructilactobacillus*, *Fructobacillus*, *Holzapfeliella*, *Nicoliella* and *Xylocopilactobacillus*, the nine genera in which at least 60% of described species have an insect or plant isolation source [3]. Genus membership was the only criterion. It holds 52 species, and every amplicon quantity rests on it.
- Tier two is species of larger mixed genera carrying an insect or plant isolation source. It holds 29 species and contributes to no amplicon denominator.

Table S1 reports the 48 species whose descriptions specify a supplement, being 39 tier-one species and nine tier-two species of *Lactobacillus*. The thirteen tier-one species not listed report no supplement, among them both species of *Holzapfeliella*. Four rules govern the record.

- A species enters the record where its description reports a supplement in the routine or the isolation medium. A supplement is recorded as required only where the description demonstrates that requirement in a complete medium, and a medium reported without such a comparison is recorded without that claim.
- Values are anchored to the nomenclatural type strain and pooled over every primary paper describing it.
- Fructophilic type is recorded in each description's own words, so the field is free text and a phrase in quotation marks is verbatim.
- Names were taken from the List on 14 April 2026 and synchronised to LactoTypeDB v1.0.0 on 13 July 2026. Departures are given in one table and applied by one script.

Each type strain's culture-collection number was resolved to a BacDive record [74] and its medium identifier to a full composition through MediaDive [75], the collection number being the key because a BacDive identifier denotes a strain rather than a species. A record was returned for 45 of the 48 type strains and 20 of the 48 resolved to a deposited medium composition. These cross-check the extracted recipe and are not reported as values.

**Reference phylogeny (Figure S1).** One type-strain genome per species was taken from LactoTypeDB v1.0.0 [4], with four outgroup type strains from three Lactobacillales families outside *Lactobacillaceae*: *Enterococcus faecalis* DSM 20478 and *Tetragenococcus halophilus* subsp. *halophilus* DSM 20339 (Enterococcaceae), *Carnobacterium divergens* DSM 20623 (Carnobacteriaceae) and *Lactococcus lactis* DSM 20481 (Streptococcaceae). Where a species offered more than one genome, a species-rank record was preferred over a nominotypical autonym and ties broken on the largest contig N50. A concatenated amino-acid alignment of Firmicutes single-copy genes was assembled with GToTree v1.8.19 run as `-H Firmicutes -N` [76], the substitution model selected with ModelTest-NG v0.1.7 [77], and the tree inferred with RAxML-NG v1.2.2 [78] under LG+I+G4+F, partitioned by gene with `--brlen scaled`, seed 42 and 100 bootstrap replicates. One ingroup genome failed the single-copy-gene filter and carries no tip. Root-to-tip distances vary several fold, so the tree is rooted on the outgroup rather than at the midpoint.

#### Amplicon dataset

**Retrieval and recruitment.** Retrieval is the acquisition of a run from the Sequence Read Archive, being the taxonomy-scoped query that defines each acquisition wave and the download that follows it. Recruitment is the permissive search of that run's chimera-free cluster seeds against the curated family reference, which decides only whether a sequence is plausibly in the family. The two act on different units. A run is retrieved and a sequence is recruited, so a recruited run, wherever the term appears below, is a run contributing at least one recruited sequence. Table S4 names the two steps Retrieval and Family recruitment in its step column.

**Metagenome pseudo-taxa.** Many amplicon runs are deposited under a metagenome label rather than a host binomial. Each of the 446 labels present in the retrieved taxonomy was curated once, and the 106 naming a source were mapped onto the isolation-source vocabulary. 34 name a plant or an invertebrate and are in scope, among them `insect gut metagenome`, `bee metagenome`, `termite gut metagenome`, `flower metagenome` and `pollen metagenome`. Ten name no host and are out: `metagenome`, `gut metagenome`, `feces metagenome`, `skin metagenome`, `oral metagenome`, `mixed sample`, `uncultured prokaryote`, `synthetic metagenome`, `fungus metagenome` and `mixed culture metagenome`. The rest resolve to a vertebrate host, an environment or a food.

A label naming no host states no habitat, so for those runs the habitat follows the host and both come from the BioSample `host` attribute. 18,320 in-scope runs take their habitat that way, every one from the `host` attribute and none from a free-text field. A run under one of the ten labels whose `host` attribute resolves to no plant or invertebrate taxon is out of scope, the Mills *et al.* dataset discussed in the main text being one. Where a label names a host and the `host` attribute names a different one, the field order below decides.

**Host resolution.** Three rules govern a host call.

- A field contributes a host only where a Linnaean name in it resolves to a single taxon carrying a family or genus within the plant or invertebrate lineages reported here. A name

resolving only above family rank contributes nothing.
- Where two names in one field resolve to different families, the run's host is left unresolved.
- A resolved host contradicting the host class the run already carries is refused.

One case departs from the stated field order. The SRA `organism` field names the taxon a run was deposited under, and is not one of the four host fields above. Where `organism` resolves to a host lineage and the `host` attribute then names that animal's own food plant, `organism` is taken, the deposited organism being the one sequenced and the `host` attribute naming its host one trophic level down. The run title is the only one of the four fields recorded per run, so it is what separates hosts within a study depositing several under one project accession.

Of the 360,693 in-scope runs, 14,732 (4.1%) carry a host resolved from a field fetched for this study, adding 17 host families and 158 genera to the analysed scope. Scored against 2,994 runs whose host family the `host` attribute already gave, with that attribute blinded, the two free-text fields called 839, left 2,155 unresolved, and reproduced the known family for 836 of those called.

**The reported unit.** No error model was fitted to any run. Three steps take the place of denoising: reads carrying a non-ACGT base were discarded, within-sample singletons were removed, and an OTU was required to recur in two independent BioProjects. Singleton removal and the fastidious graft both merge or drop and never create, so every OTU count is a lower bound. A merge spans at most two nucleotides, being 0.7% of the 292-base median V4 window and 0.4% of the 465-base V3-V4 window. Swarm was preferred to denoising on an independent benchmark of 1,800 sequence-variant tables from high-diversity environmental V3-V4 samples, in which Swarm was the least variable between replicates and DADA2 the most [33].

**Read paths.** Three paths reach quality filtering, measured over the 298,329 in-scope runs that recruited successfully and carry a per-run merge record.

- Merged reads, where a paired run merges at a rate of at least one in five: 85.1% of runs, mean identity to the nearest reference 83.5%.
- The deposited read alone, where a run carries no second read: 13.3%.
- The forward read alone, where a paired run merges below that rate: 1.6% of runs, mean identity 85.0%, and 2.6% of invertebrate runs against 1.5% of plant runs.

A shorter read carries fewer discriminating positions, so the fallback biases identity upward and cannot make a described lineage appear undescribed.

**Denominators.** 5,129 BioProjects contributed at least one recruited in-scope OTU and 3,344 carried a confirmed family detection. In 60 of those every detection resolved no deeper than family, leaving the 3,284 the main text reports (Table S2).

**Scope and losses.** Foods and environmental samples were retrieved alongside plant and invertebrate hosts and removed at one declared step (Table S2), along with vertebrate-derived, raw dairy and raw meat runs. The recurrence requirement is re-applied within the narrowed scope, removing 14,515 of the 19,256 undescribed OTUs detected there, so no reported lineage rests on an out-of-scope detection. In 104 BioProjects every run returned a filter error and contributed no OTU, so those studies enter no denominator. In a further 171 of the 8,873 BioProjects carrying a family-positive run, only some runs were lost; study size is understated for those, while sequencing depth, a median over recruited runs, is not. Neither is a reported quantity.

**Primers.** Primers were not trimmed and every run was treated the same way, SRA metadata recording a primer sequence for 0.8% of them. Retained primer bases are shared between queries but not between query and reference, so identity to the nearest type strain is biased upward and the compatible set enlarged, leaving a reported rank shallower and the undescribed class a lower bound.

**From pool to classified corpus.** Recruited reads were pooled and dereplicated across runs. Requiring occurrence in at least two independent BioProjects reduced 92,814,155 unique sequences to 5,999,101, which is the corpus-wide artefact control. A fifth acquisition wave, a sweep of fermented foods, was withdrawn before classification because its free-text scope expression admitted a majority of off-target vertebrate material, and the recurrence requirement was then re-applied within waves one to four, which left 5,698,757 OTUs for the classifier. A sequence dropped at that step is one whose two supporting BioProjects were not both in waves one to four. That set was reduced to family candidates by a permissive BLAST screen against the curated family reference at the same 80% threshold [79]. It is a computational filter and not a taxonomic one, family membership being decided by the first classification stage, and it excludes no OTU that recruitment retained except through its requirement that an alignment cover at least 55% of the query.

Amplicon region was derived from each OTU's own sequence, run metadata reporting a region for 1.1% of runs. Each OTU was aligned to the *Escherichia coli* 16S rRNA gene (GenBank J01859.1) in the conventional numbering [32] and its span binned into one of nine regions bounded by the *E. coli* coordinates of published primer pairs [80–82], one reverse-primer coordinate corrected to its measured binding site. The frame is re-checked on every run against the primer sequences, so a mis-specified frame halts the build.

#### Classification

The three stages run in the order LactoTypeDB recommends. Family membership is established before any species-level method is run, and that method's output is reported as a set. Seven genera of the family carry no label in SILVA 138.2 and take the name of their nearest labelled relative, so the genus label from stage 1 is read for family membership alone. Orientation was normalised to the plus strand between stages 1 and 2, without which a minus-strand OTU is ruled out of every species through search failure. Compatible sets were resolved to canonical species before their size was read, LactoTypeDB holding 460 records for 434 species.

**The species rule-out.** Unassigner was run with `--threshold`, `--soft_threshold` and `--ref_mismatch_positions` all unset, which is its hard default, so the species boundary is 97.5% identity over the full-length gene. For each OTU and type strain the aligned window was trimmed of end gaps, and its mismatches and matches, each incremented by a half, parameterised a beta-binomial posterior on the per-position mismatch rate. With no reference mismatch database loaded, the rate outside the window equals the rate inside it, which is the constant-rate model. That posterior was applied to the type strain's positions outside the window, returning the probability that identity over the whole gene is at least 97.5%. A species was ruled out where that probability fell to 0.5 or below, which is the cutoff the method defines [17]. The reference is the type strain of each described species and carries no environmental sequence, so a species can be ruled out only against a nomenclatural type.

Seven described species have no admissible full-length type-strain sequence. LactoTypeDB deposits a per-region insert for each, located in the type genome with barrnap [83], five of which yield a complete segment. Survivors were screened against the insert for their own region and a recovered OTU carried forward marked as resting on it. Where LactoTypeDB cannot represent a described species at an OTU's own region, that OTU is placed by the ordinary compatible-set rule and marked as such. An undescribed call is a candidate taxon awaiting the genomic confirmation required by the minimal standards for the use of genome sequences in prokaryote taxonomy [84], which this study does not provide.

All three identity thresholds were derived from full-length sequences and are applied here to region fragments, so the outcome profile is reported per region. Their re-derivation over 19,556 type strains was likewise restricted to sequences of 1,100 to 1,750 bases, whereas the windows used here are 292 bases for V4 and 465 for V3-V4. Once percentile bands replace point thresholds, adjacent ranks overlap, so an identity in the low nineties is compatible with both a new genus and a new family [85]. Identity is reported as a continuous quantity for that reason, and the 94.5% mark is drawn on the figures as a published reference point rather than a decision rule.

Of the 1,754 undescribed OTUs left unassigned at genus, 827 came from a region with no deposited collapse groups, 712 of them V5-V6, and 927 from a neighbourhood spanning more than one genus. Counted as detections, plant rows are left unassigned more often than invertebrate ones, on 3,056 of 4,870 undescribed detections against 1,398 of 6,537.

#### Controls

**Reagent and carryover screen.** Positive controls and mock communities were excluded, a mock community containing lactic acid bacteria by construction. Two panels are reported, all negative controls and those from BioProjects whose own samples are in the analysed frame, and only the second speaks to the studies analysed here. Reagent background was separated from plate carryover by the number of distinct BioProjects a genus's control detections span.

**Recovery of a published lineage.** The front end and the family-membership test were run over the sixteen runs of BioProject PRJNA925568 from brood-cell provisions of the cellophane bee *Ptiloglossa arizonensis*, a study reporting several species and strains of *Apilactobacillus* [54]. The criterion, set before the test, was that at least one run recruit *Lactobacillaceae* whose dominant best-hit genus is *Apilactobacillus* at size two or more. It was met.

**Leak-through of off-target input.** Nine full-length out-of-family 16S rRNA gene sequences were passed through 80% recruitment and then the family-membership test. The three from other phyla reach 75.7 to 76.5% identity to their nearest family member and are not recruited. The six from other families of the Bacillota and the Lactobacillales reach 85.3 to 91.0%, are recruited, and are then refused. Recruitment is permissive, and family membership is the strict test.

**Hold-out tests on the family-membership test.**

- One Lactobacillales sibling family withheld: 3 of 87 of its reads falsely called in-family.
- All five siblings withheld together: 12 of 92.
- Three families from outside the order: none of 593 reads called in-family.
- Six genera of this family dropped: family rank retained for 3,400 of 3,408 of their reads, while 629 take a wrong genus label at a bootstrap of 80 or more.
- 5,000 reads of known family membership: 4,996 called in-family.
- 564 out-of-family reads in a fourth test: none was classified at family rank.

The wrong-genus rate is why stage 1 is read for family membership alone.

**Reference completeness.** The recruitment reference and the classification reference are two different sequence sets, built from different sources. The first is a curated family reference of type-strain 16S rRNA gene records from GenBank and RefSeq, searched at 80% identity; the classification reference is LactoTypeDB v1.0.0, derived from type-strain genomes, against which every rank call is made [4]. Completeness matters to the second and not to the first, because recruitment decides only whether a read is plausibly in the family. The recruitment reference carries 473 family sequences covering 436 of the 442 species on the list it was compiled from, so 36 of the 37 genera are represented, and it is searched at 80% identity, so a member is recruited by its family neighbours rather than by its own species. That 442 is the working list's own species count. It is not the family's nomenclatural total, which the Introduction gives, and the two were taken from different sources at different dates. *Daquilactobacillus* carries no sequence in it and was recruited and named in 37 BioProjects, so a gap in the recruitment reference need not cost a detection. Under leave-one-out every sequence retains a non-self match, the median nearest-neighbour identity is 99.3%, and one sequence of 473 falls below the 80% threshold.

#### Statistical analysis

**De-duplication.** One dataset can hold two BioProject accessions, each archive issuing its own and nothing in the metadata joining them, so counting both as independent studies would be pseudoreplication. Runs deposited more than once were identified by an order-independent checksum over each run's whole OTU and abundance vector, confirmed against the archives' own per-run base counts, and the duplicates removed. Neither this step nor the curated whole-project collapse separates sibling accessions deposited by one laboratory from a single campaign, so a repeatedly sampled host can carry more accessions than independent studies.

**Independence of units.** One BioProject contributes at most one unit per host node. Within any single pooled fit between 86.5% and 100% of BioProjects contribute exactly one unit, so no estimate rests on repeated observations of a single study. Of the contrasts reported in the main text, only *Apis* against *Melipona* shares a BioProject between its two nodes, and it shares one.

**Estimator and covariates.** Firth's penalised estimator returns a finite estimate where a host node is all-positive or all-negative for a genus, which an unpenalised fit does not. The effort covariates are carried so that a difference between hosts is not a difference in sampling effort. The two host classes differ in amplicon region, extraction protocol and study design as well as in host, which is why a host-class contrast is read only against the within-region refit. Too few BioProjects are available to refit the host-order or the undescribed-rate contrasts the same way.

**Model fitting.** Each node's estimate and interval were taken from a refit with that node at the reference level, with the FLIC correction applied to both bounds. Point estimates came from `brglm2` and intervals from `logistf`, the two agreeing to a median absolute log discrepancy of 0.0004 in the odds ratio over all 2,525 contrasts. An unconverged fit was excluded. Contrasts were tested by penalised likelihood ratio, and raw prevalence accompanies every model estimate. Convergence handling and the implementation cross-check are in the software deposit. A genus's detected breadth scales with the number of species LactoTypeDB carries for it, which is one number per genus, so it cancels when the genus is held fixed and the host varied. Host nodes nest, so a host family and one of its own genera in a single fit would put two levels on the same observations; each fit covers host nodes of one rank, and every rank of every host class was fitted and deposited whether or not it is reported.

A unit whose only in-family evidence was ambiguous was excluded from every estimate, and the excluded count is reported beside every denominator. Ambiguous, unassigned and unresolved are distinct throughout. Ambiguous means an OTU resolved no deeper than family; an unassigned undescribed OTU has a nearest described relative that resolves to no single genus at that OTU's own amplicon region; a run is unresolved when no metadata field gives it a host.

**The two compositional quantities.** Lineage composition is the share of a host node's genus-resolved OTUs going to each genus, counting distinct lineages rather than abundance. Both quantities are closed, so one genus's share moves when another dominates, and each is read across the set of genera rather than between host nodes. An OTU enters the composition only where it recurs in two independent in-scope BioProjects, whereas the outcome profile's denominator is every OTU a host recruited; both denominators are stated wherever the composition is drawn.

Read share is the share of a study's family reads and is reported within single BioProjects only, where amplicon region, extraction, kit, run and site are constant (Figure S2). It is not pooled across studies, because primer choice alters recovered relative abundance [86], extraction protocol is a leading source of technical variance [87], and differences between handling laboratories exceeded the effect of specimen type [88]. A genus's *rrn* copy number scales its read share by the same factor everywhere, so it cancels across hosts and remains between two genera within a host. Copy number varies across the family [89] and correcting for it remains unsolved [90], so no correction was applied. A composition resting on fewer than 100 family reads is marked censored.

**Identity distributions.** Identity to the nearest type strain occupies a 0.1-point grid with a point mass at exactly 100, so it was drawn as a mass function with one bar per grid value and no boundary correction. It is reported as percent identity throughout.

#### Reproducibility

Every step downstream of retrieval is deterministic: iteration is sorted before any output is written, no wall-clock value reaches an artefact, seeds are fixed and both interpreters are pinned. Each deposited figure and table records the SHA-256 of every input it read, and the figure and table layer regenerates byte-identically from the deposited data.

Retrieval and the per-run front end ran in two compute environments. Acquisition waves one and two ran on Amazon Web Services EC2 spot instances, whose local scratch is ephemeral and whose capacity is pre-emptible, and waves three and four on a single local workstation. The filter errors counted in Table S2 arose almost entirely on the first, and the per-environment accounting is in the software deposit.

---

#### SUPPLEMENTARY REFERENCES

The primary species descriptions cited by Supplementary Table S1. These works are cited in the supplementary material only and are numbered separately from the article's reference list. A work cited in both carries a number in each.


1. Alcántara C, Peirotén Á, Ramón-Nuñez LA et al. Nicoliella lavandulae sp. nov., a novel fructophilic Nicoliella species isolated from flowers of Lavandula angustifolia. *Int J Syst Evol Microbiol* 2024;74:006497. https://doi.org/10.1099/ijsem.0.006497

2. Antunes A, Rainey FA, Nobre MF et al. Leuconostoc ficulneum sp. nov., a novel lactic acid bacterium isolated from a ripe fig, and reclassification of Lactobacillus fructosus as Leuconostoc fructosum comb. nov. *Int J Syst Evol Microbiol* 2002;52:647–55. https://doi.org/10.1099/ijs.0.02004-0

3. Botero J, Peeters C, De Canck E et al. Eupransor demetentiae gen. nov., sp. nov., a novel fructophilic lactic acid bacterium from bumble bees. *Int J Syst Evol Microbiol* 2024;74:006409. https://doi.org/10.1099/ijsem.0.006409

4. Botero J, Peeters C, De Canck E et al. A comparative genomic analysis of Fructobacillus evanidus sp. nov. from bumble bees. *Syst Appl Microbiol* 2024;47:126505. https://doi.org/10.1016/j.syapm.2024.126505

5. Chambel L, Chelo IM, Zé-Zé L et al. Leuconostoc pseudoficulneum sp. nov., isolated from a ripe fig. *Int J Syst Evol Microbiol* 2006;56:1375–81. https://doi.org/10.1099/ijs.0.64054-0

6. Charlton DB, Nelson ME, Werkman CH. Physiology of Lactobacillus fructivorans sp. nov., isolated from spoiled salad dressing. *Iowa State Coll J Sci* 1934;9:1–11.

7. Chen YS, Wang LT, Lin ST et al. Fructobacillus apis sp. nov., isolated from the gut of honeybee (Apis mellifera). *Int J Syst Evol Microbiol* 2022;72:005613. https://doi.org/10.1099/ijsem.0.005613

8. Chiou TY, Suda W, Oshima K et al. Lactobacillus kosoi sp. nov., a fructophilic species isolated from kôso, a Japanese sugar-vegetable fermented beverage. *Antonie van Leeuwenhoek* 2018;111:1149–56. https://doi.org/10.1007/s10482-018-1019-7

9. Deshmukh UB, Oren A. Proposal of Christiangramia gen. nov., Neomelitea gen. nov. and Nicoliella gen. nov. as replacement names for the illegitimate prokaryotic generic names Gramella Nedashkovskaya et al. 2005, Melitea Urios et al. 2008 and Nicolia Oliphant et al. 2022, respectively. *Int J Syst Evol Microbiol* 2023;73:005806. https://doi.org/10.1099/ijsem.0.005806

10. Edwards CG, Haag KM, Collins MD et al. Lactobacillus kunkeei sp. nov.: a spoilage organism associated with grape juice fermentations. *J Appl Microbiol* 1998;84:698–702. https://doi.org/10.1046/j.1365-2672.1998.00399.x

11. Endo A, Okada S. Reclassification of the genus Leuconostoc and proposals of Fructobacillus fructosus gen. nov., comb. nov., Fructobacillus durionis comb. nov., Fructobacillus ficulneus comb. nov. and Fructobacillus pseudoficulneus comb. nov. *Int J Syst Evol Microbiol* 2008;58:2195–205. https://doi.org/10.1099/ijs.0.65609-0

12. Endo A, Futagawa-Endo Y, Sakamoto M et al. Lactobacillus florum sp. nov., a fructophilic species isolated from flowers. *Int J Syst Evol Microbiol* 2010;60:2478–82. https://doi.org/10.1099/ijs.0.019067-0

13. Endo A, Irisawa T, Futagawa-Endo Y et al. Fructobacillus tropaeoli sp. nov., a fructophilic lactic acid bacterium isolated from a flower. *Int J Syst Evol Microbiol* 2011;61:898–902. https://doi.org/10.1099/ijs.0.023838-0

14. Gallus MK, Beer I, Ivleva NP et al. Fructobacillus cardui sp. nov., isolated from a thistle (Carduus nutans) flower. *Int J Syst Evol Microbiol* 2022;72:005553. https://doi.org/10.1099/ijsem.0.005553

15. Jiang CS, Gu CT. Lactobacillus juensis sp. nov. and Lactobacillus rizhaonensis sp. nov., isolated from the gut of honeybee (Apis mellifera). *Int J Syst Evol Microbiol* 2024;74:006285. https://doi.org/10.1099/ijsem.0.006285

16. Kanaya R, Maeno S, Hisatomi A et al. Apilactobacillus pseudapinorum sp. nov., isolated from polyfloral honey. *Int J Syst Evol Microbiol* 2026;76:007027. https://doi.org/10.1099/ijsem.0.007027

17. Kawasaki S, Ozawa K, Mori T et al. Symbiosis of carpenter bees with uncharacterized lactic acid bacteria showing NAD auxotrophy. *Microbiol Spectr* 2023;11:e00782–23. https://doi.org/10.1128/spectrum.00782-23

18. Kitahara K, Kaneko T, Goto O. Taxonomic studies on the hiochi-bacteria, specific saprophytes of sake. *J Gen Appl Microbiol* 1957;3:111–20. https://doi.org/10.2323/jgam.3.111

19. Kline L, Sugihara TF. Microorganisms of the San Francisco sour dough bread process. *Appl Microbiol* 1971;21:459–65. https://doi.org/10.1128/am.21.3.459-465.1971

20. Kodama R. Studies on the nutrition of lactic acid bacteria. Part IV. Lactobacillus fructosus nov. sp., a new species of lactic acid bacteria. *Nippon Nogeikagaku Kaishi* 1956;30:705–9.

21. Leisner JJ, Vancanneyt M, Van der Meulen R et al. Leuconostoc durionis sp. nov., a heterofermenter with no detectable gas production from glucose. *Int J Syst Evol Microbiol* 2005;55:1267–70. https://doi.org/10.1099/ijs.0.63434-0

22. Li TT, Gu CT. Lactobacillus huangpiensis sp. nov. and Lactobacillus laiwuensis sp. nov., isolated from the gut of honeybee (Apis mellifera). *Int J Syst Evol Microbiol* 2022;72:005237. https://doi.org/10.1099/ijsem.0.005237

23. Lin ST, Guu JR, Wang HM et al. Fructobacillus papyriferae sp. nov., Fructobacillus papyrifericola sp. nov., Fructobacillus broussonetiae sp. nov. and Fructobacillus parabroussonetiae sp. nov., isolated from paper mulberry in Taiwan. *Int J Syst Evol Microbiol* 2022;72:005235. https://doi.org/10.1099/ijsem.0.005235

24. McFrederick QS, Vuong HQ, Rothman JA. Lactobacillus micheneri sp. nov., Lactobacillus timberlakei sp. nov. and Lactobacillus quenuiae sp. nov., lactic acid bacteria isolated from wild bees and flowers. *Int J Syst Evol Microbiol* 2018;68:1879–84. https://doi.org/10.1099/ijsem.0.002758

25. Oliphant SA, Watson-Haigh NS, Sumby KM et al. Apilactobacillus apisilvae sp. nov., Nicolia spurrieriana gen. nov. sp. nov., Bombilactobacillus folatiphilus sp. nov. and Bombilactobacillus thymidiniphilus sp. nov., four new lactic acid bacterial isolates from stingless bees Tetragonula carbonaria and Austroplebeia australis. *Int J Syst Evol Microbiol* 2022;72:005588. https://doi.org/10.1099/ijsem.0.005588

26. Oliphant SA, Watson-Haigh NS, Sumby KM et al. Fructilactobacillus cliffordii sp. nov., Fructilactobacillus hinvesii sp. nov., Fructilactobacillus myrtifloralis sp. nov., Fructilactobacillus carniphilus sp. nov. and Fructobacillus americanaquae sp. nov., five novel lactic acid bacteria isolated from insects or flowers of Kangaroo Island, South Australia. *Int J Syst Evol Microbiol* 2023;73:005730. https://doi.org/10.1099/ijsem.0.005730

27. Olofsson TC, Alsterfjord M, Nilson B et al. Lactobacillus apinorum sp. nov., Lactobacillus mellifer sp. nov., Lactobacillus mellis sp. nov., Lactobacillus melliventris sp. nov., Lactobacillus kimbladii sp. nov., Lactobacillus helsingborgensis sp. nov. and Lactobacillus kullabergensis sp. nov., isolated from the honey stomach of the honeybee Apis mellifera. *Int J Syst Evol Microbiol* 2014;64:3109–19. https://doi.org/10.1099/ijs.0.059600-0

28. Pham VD, Gänzle MG. Fructilactobacillus frigidiflavus sp. nov., a pigmented species, and Levilactobacillus lettrarii sp. nov., a propionate-producing species isolated from sourdough.

*Int J Syst Evol Microbiol* 2025;75:006726. https://doi.org/10.1099/ijsem.0.006726

29. Techo S, Miyashita M, Shibata C et al. Lactobacillus ixorae sp. nov., isolated from a flower (West-Indian jasmine). *Int J Syst Evol Microbiol* 2016;66:5500–5. https://doi.org/10.1099/ijsem.0.001547

30. Tsai LJ, Chen PC, Lan WT et al. Apilactobacillus intestinapis sp. nov., isolated from the honeybee. *Int J Syst Evol Microbiol* 2025;75:006971. https://doi.org/10.1099/ijsem.0.006971

31. Wang C, Huang Y, Li L et al. Lactobacillus panisapium sp. nov., from honeybee Apis cerana bee bread. *Int J Syst Evol Microbiol* 2018;68:703–8. https://doi.org/10.1099/ijsem.0.002538

32. Weiss N, Schillinger U. Lactobacillus sanfrancisco sp. nov., nom. rev. *Syst Appl Microbiol* 1984;5:230–2. https://doi.org/10.1016/s0723-2020(84)80024-7

33. Zheng J, Wittouck S, Salvetti E et al. A taxonomic note on the genus Lactobacillus: Description of 23 novel genera, emended description of the genus Lactobacillus Beijerinck 1901, and union of Lactobacillaceae and Leuconostocaceae. *Int J Syst Evol Microbiol* 2020;70:2782–858. https://doi.org/10.1099/ijsem.0.004107

### Supplementary Figures

**Fig. S1. Maximum-likelihood core-genome phylogeny of 412 *Lactobacillaceae* type-strain genomes**, one per validly published species with a clean type-strain genome, inferred from a concatenated amino-acid alignment of 119 single-copy core genes (21,859 sites) under LG+I+G4+F, selected by ModelTest-NG v0.1.7, in RAxML-NG v1.2.2 with 100 Felsenstein bootstrap replicates, seed 42 and scaled branch lengths. The alignment was assembled with GToTree v1.8.19 against the Firmicutes single-copy gene set. Each tip gives the species name in italics followed by its genome assembly accession. A dot marks each of the 385 internal splits, of 413, with bootstrap support of at least 70; support below that is not marked. The tree is rooted on an outgroup of four type strains from three Lactobacillales families outside *Lactobacillaceae*, labelled in grey: *Enterococcus faecalis* DSM 20478 and *Tetragenococcus halophilus* subsp. *halophilus* DSM 20339 (Enterococcaceae), *Carnobacterium divergens* DSM 20623 (Carnobacteriaceae) and *Lactococcus lactis* DSM 20481 (Streptococcaceae). Root-to-tip distances vary several fold across this cohort and the two outgroup-free rooting criteria disagreed about one genus, which makes midpoint rooting unreliable here. One cohort genome did not pass the single-copy-gene filter and has no tip on this tree: *Oenococcus alcoholitolerans* (GCA_000769695.1). The scale bar is in mean amino-acid substitutions per site.

One name departs from the current LPSN correct name: GCF_900636965.1 reported as *Lacticaseibacillus rhamnosus* rather than *Lactobacillus rhamnosus*. The departure is applied at the reporting layer only, tip accessions are unchanged from the alignment, and with it applied all 37 genera are monophyletic on this tree. Isolation source and cultivation conditions are not annotated here; they are given in the text, per species, each value cited to the original description.

**Alt text:** A rooted maximum-likelihood phylogenetic tree drawn as a vertical cladogram with 412 Lactobacillaceae tips, one per described species, each labelled with its species name and genome assembly accession. The four outgroup tips from three other Lactobacillales families are labelled in grey at the base and the tree is rooted on them. Branch lengths are drawn to scale in amino-acid substitutions per site, with a scale bar. Species of one genus fall together in contiguous, well-separated clades across the tree. A filled dot on an internal node marks bootstrap support of at least 70; 385 of the 413 internal splits carry one, so the great majority of the branching order is strongly supported.

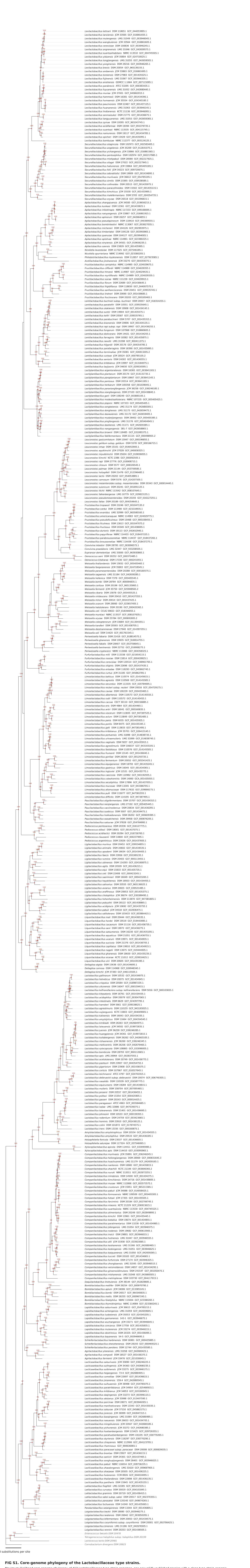

**FIG S1. Core-genome phylogeny of the Lactobacillaceae type strains.**

Maximum-likelihood core-genome phylogeny of 412 Lactobacillaceae type-strain genomes, one per validly published species with a clean type-strain genome, inferred from a concatenated amino-acid alignment of 119 single-copy core genes (21,859 sites) under LG+I+G4+F in RAxML-NG v1.2.2 with 100 Felsenstein bootstrap replicates, seed 42 and scaled branch lengths. Each tip gives the species name in italics, then the type-strain designation and the genome assembly accession in roman. A red dot marks each of the 385 internal splits, of 413, with bootstrap support of at least 70 per cent, and the value is printed beside it; support below that is not marked. The tree is rooted on an outgroup of four type strains from three Lactobacillales families outside Lactobacillaceae, labelled in grey. Root-to-tip distances vary several fold across this cohort, which makes midpoint rooting unreliable. One cohort genome did not pass the single-copy-gene filter and has no tip on this tree: Oenococcus alcoholitolerans (GCA_000769695.1). The scale bar is in mean amino-acid substitutions per site.

**Fig. S2. Within-study composition of the *Lactobacillaceae* across hymenopteran host genera.** Each panel is one BioProject, so the amplicon region, the extraction and library kit, the sequencing run, the collection site and the reference are held constant and a difference between host genera within a panel is not a batch effect. 6 BioProjects and 31 host-genus rows are drawn, each row a host genus with at least 4 runs in that project; 15 rows fall below that threshold and are not drawn. Amplicon region per project: PRJNA1150198 V5-V6; PRJNA309422 V4; PRJNA429121 V4; PRJNA749807 V5-V6; PRJNA798723 V4; PRJNA862864 V4.

The composition axis is each named genus's share of the family's reads, with the ambiguous and undescribed fractions shown as hatched bands rather than dropped. An ambiguous band wide enough to carry text is labelled with its compatible set, naming as many genera as fit within the band and counting the rest, and giving the top set's share of the band where no one set holds most of it. The right-hand axes give the family's share of all reads and the absolute family read count, with the censoring threshold marked.

Panels are read down a column and not across a row. *rrn* copy number is a property of a genus, so it cancels when one genus is compared between hosts and does not cancel when two genera are compared within a host. This is the only contrast in which those variables are controlled rather than adjusted for, and it is an existence demonstration rather than a corpus-scale estimate; no rate in the main text is computed from it.

**Alt text:** A stack of 6 panels, one per BioProject, holding 31 horizontal bars in all, each bar a host genus within that project. Each bar is a 100 per cent stacked composition of that host's Lactobacillaceae reads, one coloured segment per named genus with the ambiguous and undescribed fractions drawn as hatched segments rather than dropped. Two axes run down the right-hand side of every panel: the family's share of all reads, and the absolute family read count on a log scale with the censoring threshold marked by a vertical rule. Because a panel holds one sequencing run, region, kit and site, the figure is read down a column, comparing one host genus with another inside a panel, and not across panels.

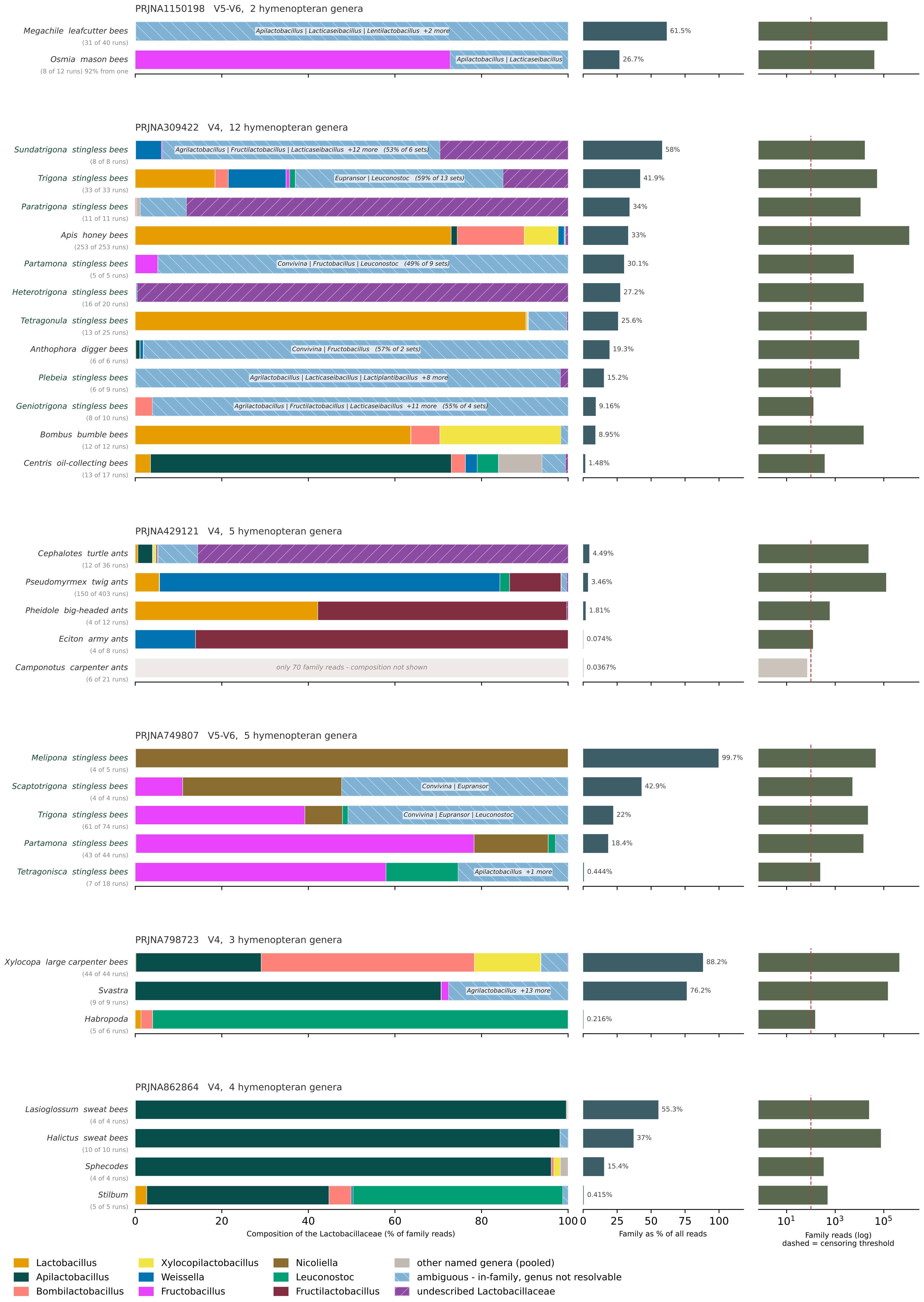
PRJNA1150198 V5-V6, 2 hymenopteran genera
Megachile leafcutter bees
(31 of 40 runs)
Apilactobacillus | Lacticaseibacillus | Lentilactobacillus +2 more
61.5%
Osmia mason bees
(8 of 12 runs) 92% from one
Apilactobacillus | Lacticaseibacillus
26.7%
PRJNA309422 V4, 12 hymenopteran genera
Sundatrigona stingless bees
(8 of 8 runs)
Agrilactobacillus | Fructilactobacillus | Lacticaseibacillus +12 more (53% of 6 sets)
58%
Trigona stingless bees
(33 of 33 runs)
Eupransor | Leuconostoc (59% of 13 sets)
41.9%
Paratrigona stingless bees
(11 of 11 runs)
34%
Apis honey bees
(253 of 253 runs)
33%
Partamona stingless bees
(5 of 5 runs)
Convivina | Fructobacillus | Leuconostoc (49% of 9 sets)
30.1%
Heterotrigona stingless bees
(16 of 20 runs)
27.2%
Tetragonula stingless bees
(13 of 25 runs)
25.6%
Anthophora digger bees
(6 of 6 runs)
Convivina | Fructobacillus (57% of 2 sets)
19.3%
Plebeia stingless bees
(6 of 9 runs)
Agrilactobacillus | Lacticaseibacillus | Lactiplantibacillus +8 more
15.2%
Geniotrigona stingless bees
(8 of 10 runs)
Agrilactobacillus | Fructilactobacillus | Lacticaseibacillus +11 more (55% of 4 sets)
9.16%
Bombus bumble bees
(12 of 12 runs)
8.95%
Centris oil-collecting bees
(13 of 17 runs)
1.48%
PRJNA429121 V4, 5 hymenopteran genera
Cephalotes turtle ants
(12 of 36 runs)
4.49%
Pseudomyrmex twig ants
(150 of 403 runs)
3.46%
Pheidole big-headed ants
(4 of 12 runs)
1.81%
Eciton army ants
(4 of 8 runs)
0.074%
Camponotus carpenter ants
(6 of 21 runs)
only 70 family reads - composition not shown
0.0367%
PRJNA749807 V5-V6, 5 hymenopteran genera
Melipona stingless bees
(4 of 5 runs)
99.7%
Scaptotrigona stingless bees
(4 of 4 runs)
Convivina | Eupransor
42.9%
Trigona stingless bees
(61 of 74 runs)
Convivina | Eupransor | Leuconostoc
22%
Partamona stingless bees
(43 of 44 runs)
18.4%
Tetragonisca stingless bees
(7 of 18 runs)
Apilactobacillus +1 more
0.444%
PRJNA798723 V4, 3 hymenopteran genera
Xylocopa large carpenter bees
(44 of 44 runs)
88.2%
Svastra
(9 of 9 runs)
Agrilactobacillus +13 more
76.2%
Habropoda
(5 of 6 runs)
0.216%
PRJNA862864 V4, 4 hymenopteran genera
Lasioglossum sweat bees
(4 of 4 runs)
55.3%
Halictus sweat bees
(10 of 10 runs)
37%
Sphecodes
(4 of 4 runs)
15.4%
Stilbum
(5 of 5 runs)
0.415%
0
20
40
60
80
100
Composition of the Lactobacillaceae (% of family reads)
0
25
50
75
100
Family as % of all reads
$10^1$
$10^3$
$10^5$
Family reads (log)
dashed = censoring threshold
Lactobacillus
Apilactobacillus
Bombilactobacillus
Xylocopilactobacillus
Weissella
Fructobacillus
Nicoliella
Leuconostoc
Fructilactobacillus
other named genera (pooled)
ambiguous - in-family, genus not resolvable
undescribed Lactobacillaceae
