## Supplementary File Summaries for "Stingless bees, turtle ants and tea plants are rich sources of undescribed *Lactobacillaceae*"

Supplementary file summaries
The two supplementary files accompanying this preprint, with their contents and formats.

Supplementary_Material.pdf (PDF)
Supplementary Methods for the cultivation record, the amplicon dataset, classification,
controls, statistical analysis and reproducibility, with its own reference list, followed
by Supplementary Figures S1 and S2 with their legends. Sixteen pages.

Supplementary_Tables.xlsx (Microsoft Excel workbook)
Supplementary Tables S1 to S16 as twenty-one sheets, one table per sheet, each carrying
its caption, with a Legends sheet listing all twenty-one.
